# Predicting and Designing therapeutics against the Nipah virus

**DOI:** 10.1101/623603

**Authors:** Neeladri Sen, Tejashree Rajaram Kanitkar, Ankit Animesh Roy, Neelesh Soni, Kaustubh Amritkar, Shreyas Supekar, Sanjana Nair, Gulzar Singh, M.S. Madhusudhan

## Abstract

Though every outbreak of the Nipah Virus has resulted in high mortality rates (>70% in Southeast Asia), there are no licensed drugs against it. In this study we have considered all 9 Nipah proteins as potential therapeutic targets and computationally identified putative peptides (against G, F, and M proteins) and small molecules inhibitors (against F, G, M, N, and P proteins). The computations include extensive homology/*ab initio* modelling, peptide design and small molecule docking. An important contribution of this study is the increased structural characterization of Nipah proteins by approximately 90% of what is deposited in the PDB. In addition, we have carried out molecular dynamics simulations on all the designed protein-peptide complexes to check for stability and to estimate binding strengths. Details, including atomic coordinates of all the proteins and their ligand bound complexes, can be accessed at http://cospi.iiserpune.ac.in/Nipah. Our strategy was to tackle the development of therapeutics on a proteome wide scale and the lead compounds identified could be attractive starting points for drug development. To this end, we have designed 4 peptide inhibitors and predicted 70 small molecules (13 with high confidence) against 3 and 5 Nipah proteins respectively. To counter the threat of drug resistance, we have analyzed the sequences of the viral strains from different outbreaks, to check whether they would be sensitive to the binding of the proposed inhibitors.

**Author Summary:** Nipah virus infections have killed 72-86% of the infected individuals in Bangladesh and India. The infections are spread via bodily secretions of bats, pigs and other infected individuals. Even though, the disease was first detected in the human population in 1998, there are no approved drugs/vaccines against it. In this study, we have tried to model the 3D structures of the Nipah virus proteins. We have then used these models to design/predict small inhibitory molecules that would bind them and prevent their function. We have also analyzed the different strains of the virus to identify conservation patterns of amino acids in the proteins, which in turn informs us of the efficacy of the drugs. The designed/docked compounds as well as the protein models are freely accessible for experimental validation and hypothesis testing.

## Introduction

The May 2018 outbreak of the Nipah Virus (NiV) in Kerala, India, claimed the lives of 21 of the 23 infected people [1]. This zoonotic pathogen was first detected to infect humans in an outbreak in Malaysia in 1998 [2]. Since then, the mortality rate, especially in the Indian subcontinent has been high with Bangladesh and India reporting 72% and 86% fatalities respectively [3–5]. Though the overall number of fatalities linked with each outbreak has never been more than 105, NiV poses a deadly threat and could potentially become pandemic [6]. Considering its high mortality and transmission rates, NiV features in the WHO R&D Blueprint list of epidemic threats that need immediate R&D action [3]. In light of this, the Coalition for Epidemic Preparedness Innovations (CEPI) has extended US$ 25 million support to Profectus BioSciences, Inc. and Emergent BioSolutions Inc. for the development of vaccines against NiV in 2018 [7].

NiV is currently classified as a Biosafety Level 4 (BSL-4) pathogen [8] with no licensed drugs or vaccines. While drugs such as ribavirin and 4’-Azidocytidine [9] have shown inhibitory action *in vitro*, their *in vivo* efficacy has not been reported. Some of these drugs target the *Paramyxoviridae* family, of which NiV is a member. The drug favipiravir [10] protects against lethal doses of NiV in hamster models and is in Phase II of clinical trials. However, *in-vitro* studies have shown the emergence of resistance to this drug among members of the influenza family [11]. A monoclonal antibody, m102.4 [12] acts against the G protein of the virus has been shown to be effective on animal models but human trials are yet to be conducted, though preliminary indications appear promising [13]. In principle, structure based rational design of therapeutics and drugs could help combat the disease and also address the concerns of drug resistance.

In this study, we have comprehensively characterized the structural proteome of NiV and explored the possibility of targeting most if not all its proteins for inhibitor/drug discovery. The NiV genome encodes six structural proteins *viz*. Glycoprotein (G), Fusion protein (F), Matrix protein (M), Nucleoprotein (N), RNA-directed RNA polymerase (L), Phosphoprotein (P) and three non-structural proteins named W, C and V [14,15]. The G protein helps in viral attachment to host cell ephrin receptors and the F protein mediates its fusion [15]. The P protein binds to the N protein and maintains it in a soluble form and increases its specificity towards viral RNA instead of non-specific cellular RNA [16]. The N-P protein complex then binds to the viral RNA forming the nucleocapsid. This nucleocapsid coated viral RNA acts as a template for viral polymerase L to replicate itself and the host machinery is then utilized to translate its proteins [15]. After replication, the M protein enables viral assembly and budding/release of new viral particles [15]. The non-structural proteins W, V, and C act against the interferon signaling to escape the host immune response [15]. All these proteins are potential targets for rational drug design.

In this study, we have used the experimentally determined structures of the NiV proteins and built models for the remaining proteins to find putative lead compounds against the virus. Four proteins (F, G, N and P proteins) have structural data available in the Protein Data Bank (PDB) [17] with varying degree of structural coverage (Table 1). Using homology based methods; we have extended the structural coverage of these proteins and built models for four of the remaining proteins using either homology modeling or threading/*ab initio* methods. We designed peptide inhibitors targeting interacting sites on G protein-human ephrin-B2 receptor, F protein trimer and M protein dimer. Binding stability of inhibitory peptides was assessed with molecular dynamics (MD) simulations. In addition, to quantify the binding affinities, binding free energies of the designed peptide inhibitors to their respective targets were also evaluated, based on configurations from MD simulations. We have predicted putative drug like molecules using molecular docking that could bind to NiV proteins. Our proposed inhibitors should potentially bind to viral proteins and hinder their function thereby preventing viral life-cycle progression. Finally, we have compared the proteomes of Malaysian, Bangladesh and Indian NiV isolates for sequence variations and mapped them onto their protein structures. This enables us to delineate the consequences (if any) of sequential variation among strains on the efficacy of proposed drugs.

**Table 1.** List of NiV proteins with their lengths, PDB codes of crystal structures, coverage of crystal structures, coverage of models, additional coverage obtained by the models and the overall sequence coverage. In cases where models have increased the coverage over existing crystal structures, the original coverage is in parentheses.

| Sr. no. | Protein | Length | X-ray structures | X-ray coverage | Model coverage | Additional coverage | Overall coverage (%) |
| --- | --- | --- | --- | --- | --- | --- | --- |
| 1 | Pre fusion F protein | 546 | 5EVM, 1WP7, 3N27 | 27 - 482 | 27 - 482 | 0 | 84 |
|  | Post fusion F protein |  | - | - | 72-418 | 347 | 64 |
| 2 | G protein | 602 | 2VSM, 2VWD, 3D11, 3D12 | 176 - 602 | 98 - 597 | 79 | 71 (84) |
| 3 | N protein | 532 | 4CO6 | 32 - 371 | 39 - 414 | 44 | 64 (72) |
| 4 | P protein | 709 | 4CO6, 4GJW | 1 - 38 | 655 - 709 | 55 | 20 (28) |
|  |  |  |  | 471 - 573 |  |  |  |
| 5 | M protein | 352 | - | - | 45 - 352 | 308 | 88 |
| 6 | L protein | 2244 | - | - | 1814 - 2024 | 210 | 9 |
| 7 | V protein | 456 | - | - | 1 - 38 | 297 | 65 |
|  |  |  |  |  | 87 - 243 |  |  |
|  |  |  |  |  | 313 - 414 |  |  |
| 8 | W protein | 450 | - | - | 1 - 38 | 266 | 59 |
|  |  |  |  |  | 87 - 243 |  |  |
|  |  |  |  |  | 321 - 391 |  |  |
| 9 | C protein | 166 | - | - | - | - | - |

## Methods

### 1. Protein structure modeling

At the time of modeling, the sequence of the Indian strain was not available and so all the modeling was carried out using the Malaysian strain (AY029768.1) [18]. From our experience, using one strain over another would only minimally affect the computed models (Refer Results Section 4 for details on sequence conservation). Monomeric structures of the proteins were built using the homology modeling pipeline ModPipe-2.2.0 [22] and their multimeric complexes were built using MODELLER v9.17 [20,21]. The templates for homology modeling were identified using both sequence-sequence and profile-sequence search methods. Profile-sequence search methods improve identification of distant homologs that have sequence identity lower than 30%. The sequence profiles of the target proteins were generated using PSI-BLAST [22] against the UniRef90 database [23] with three iterations and an e-value threshold of 0.001. Models were built with dynamic Coulomb (electrostatic) restraints and were subjected to the ‘very slow’ mode of refinement with two rounds of optimization. The quality of the generated models was assessed using the Modpipe quality score, GA341, Discrete Optimized Protein Energy (DOPE) and Normalized DOPE scores [24]. Protein structure models were retained for further analyses only if they had a Normalized DOPE score less than or equal to zero.

Protein domains/regions that could not be reliably modeled by MODELLER (either greater than zero Normalized DOPE score or with less than 50% structural coverage) were rebuilt using meta-threading and *ab initio* methods on the I-TASSER web server [25]. Models built using I-TASSER were assessed with Normalized DOPE scores along with their C-scores, predicted TM scores and RMSD scores provided by the webserver [25].

### 2. Modeling inhibitory peptides against NiV proteins and assessing their stability

One peptide inhibitor was computationally designed against each of the F and M proteins while 2 inhibitors were designed against the G protein. Details of the procedure are stated in the results section. MD simulations were carried out in triplicates for all four predicted protein-peptide inhibitor complexes. The simulations were carried out using Gromacs [26,27] with the Amber99SB-ILDN force field [28]. A cubic water box whose sides were at a minimum distance of 1.2 nm from any protein atom was used for solvating each of the systems. Sodium or chloride counter ions were added to achieve charge neutrality. Electrostatic interactions were treated using the particle mesh Ewald sum method [29] and LINCS [30] was used to constrain hydrogen bond lengths. A time step of 2 fs was used for the integration. The whole system was minimized for 5000 steps or till the maximum force was less than 1000 kJ/mol/nm. The system was then heated to 300K in an NVT ensemble simulation for 100 ps using a Berendsen thermostat [31]. The pressure was stabilized in an NPT ensemble simulation for 100 ps using a Parrinello-Rahman barostat [32]. The system was simulated for a maximum of 100 ns and structures were stored after every 10 ps. The temperature, potential energy and kinetic energy were monitored during the simulation to check for anomalies.

Free energy of binding of the putative peptide inhibitors provides an important quantitative description of its efficacy. In this study, the extensive MD simulations of protein-peptide complexes were post-processed to obtain binding free energy estimates using the molecular mechanics Poisson-Boltzmann surface area (MM/PBSA) approach [33,34]. The MM/PBSA method employs an implicit solvation model to estimate the free energy of binding by evaluating ensemble averaged classical interaction energies (MM) and continuum solvation free energies (PBSA) of the protein-ligand complex conformations from the MD trajectories. Snapshots of protein-peptide complexes were obtained at every 100 ps from the last 50 ns of the MD trajectories, thus totaling 500 snapshots. The last 50 ns were selected for MM/PBSA treatment to ensure sampling of equilibrium conformations for appropriate MM/PBSA energy evaluations (Supporting Figures 2, 3, 4 and 5 for RMSD and distance between the center of peptide and protein). The MD snapshots were energy minimized for 2000 steps before evaluation of interaction and solvation free energies. The protein and solvent were modeled with dielectric constants of ε=2 and ε=80, respectively. APBS suite [35] and GMXPBSA [36] were used for implicit solvent calculations. In this study, we attempted to calculate the entropic estimate of binding using the interaction entropy formalism [37]. However, converged entropic values with reasonable error estimates for protein-peptide trajectories could not be obtained, which is often the case when evaluating entropic contributions from molecular simulations. We, therefore neglected entropic contributions to the binding free energies, which is a common practice in MM/PBSA literature [38]. The enthalpies of binding obtained from MM/PBSA calculations are reported as binding energies for the protein-peptide complexes.

### 3. Prediction of putative small molecules that can bind to NiV proteins

Docking was used to identify putative small molecules that can potentially bind and inhibit the activities of the NiV proteins. In this exercise, NiV proteins (G, N, F, P and M proteins) that had structures or models with reliable quality (Normalized DOPE <= 0) and high coverage (> 80%) were used as targets for a ligand screening. The screening library consisted of the 70% non-redundant set of clean drug like molecules of the ZINC database [39,40]. The binding pockets for docking on the targets were predicted using the DEPTH server [41,42]. The parameters of DEPTH included a minimum number of neighborhood waters set to 4 and the probability threshold for binding site of 0.8. Evolutionary information was also included by the server in cavity prediction [43]. Docking was performed using Autodock4 [44], and DOCK6 [45,46].

The target proteins were prepared for docking by Autodock4, by adding missing polar hydrogen atoms and Gasteiger charges. The ligand docking site, marked by affinity grids were generated using the Autogrid module of Autodock. The center of the grid, number of grid points in X, Y, Z direction and separation of grid points were chosen based on the predicted binding pockets using the ADT viewer from MGL tools [44]. The number of Genetic Algorithm runs was set to 20. The final energies reported by Autodock4 were used for evaluation and selection of the putative leads.

The target proteins were prepared for docking by DOCK6.8 using Dock Prep tool [45] from Chimera [47]. Missing hydrogen atoms were added to the target proteins. Charges on atoms of the protein were determined using AMBER. Molecular surface of the target was generated using the DMS tool from Chimera. The sphgen program from DOCK6.8 was used to generate spheres from the molecular surface. The cluster of spheres were selected according to the binding sites predicted by DEPTH. The grid box and grid were created by showbox and grid programs respectively. Flexible ligand docking was performed using DOCK6. The final energies reported by DOCK6.8 were used for evaluation and selection of the putative leads.

### 4. Mapping strain variants onto structure

Protein sequences of 15 different NiV isolates, 7 from Malaysia (AY029768.1,A J564621.1, AJ627196.1, AY029767.1, AJ564622.1, AJ564623.1, AF212302.2) [18], 3 from Bangladesh (AY988601.1, JN808857.1, AY988601.1) [48] and 5 from India (MH523641.1, MH523642.1, MH396625.1, MH523640.1, FJ513078.1)[49] were retrieved from their translated genomes deposited in the NCBI nucleotide database [50] and were used to identify sequence variations in proteins. We also verified that the translated protein sequences of the Malaysian strain matched with those of the protein sequences deposited in SwissProt [51]. Multiple sequence alignments of the sequences obtained from the 15 isolates were performed with MUSCLE [52]. Positions with amino acid variations were mapped onto the structures. Amino acid variations within 5 at inhibitor binding sites were identified.

## Results

### 1. Structural coverage of the NiV proteome

#### Homology modeling the Nipah proteome

In this study, we first focused on characterizing the structures of the NiV proteins. Partial structures for 4 (F, G, N and P protein) of the 9 NiV proteins are available in the PDB (Table 1). Computationally, we attempted to extend the structural coverage of these 4 proteins and to build models for the remaining 5 proteins using homology modeling (with MODELLER), *ab initio* modeling and threading (with I-TASSER). Model accuracies were carefully scrutinized before attempts to design/find inhibitors against all possible proteins in the proteome. In this section, we only present the results of homology modeling as all models built using I-TASSER resulted in structures that were not favorably assessed (Normalized DOPE > 0) (Supporting Table 1)

Multiple models were constructed for each of the proteins using all available templates. All proteins, except C, had at least one model with a normalized DOPE score of less than or equal to zero. All models built for proteins with existing X-ray structure conferred additional sequence coverage except for the F protein (Table 1). The structural coverage of the N, P and G proteins increased by 8-13% after modeling (Table 1). Overall, we increased the structural coverage of the NiV proteome by 90%, from ~23% (1364 residues) to ~43% (2623 residues).

#### Modeling the post-fusion F protein

For NiV to enter a host’s cell, its G protein attaches to the host ephrin receptor and the F protein is instrumental in fusing the viral envelope with the host cell membrane [53]. The F protein undergoes a conformational change from the pre-fusion to the post-fusion state triggered by the binding of the G protein to the ephrin receptor. These conformational changes are characteristic of class I viral fusion proteins [53–58]. The structure of only the pre-fusion state of the NiV F protein has been determined experimentally (PDB ID: 5EVM). We modeled the post-fusion state using the structure of the human Parainfluenza Virus 3 (PDB ID: 1ZTM) as a template since it is also a class I fusion protein. Though the NiV and human parainfluenza virus fusion proteins are only 26.4 % identical in sequence, their pre-fusion conformations take on similar folds with a structure overlap of 67% and an RMSD of 0.2 nm (as calculated using CLICK [59]). The rationale for modelling the post-fusion state of NiV using the Parainfluenza virus template is further corroborated by reports in literature of the common mode of conformational change in post-fusion states of class I viral fusion proteins [53,60–64] despite their low sequence identity (Supporting Figure 1) leading to the formation of 6 helix bundle. The target-template alignment was done using CLUSTALW-1.7, and the model was constructed using MODELLER v9.17. It has previously been shown that Hendra virus (HeV) and NiV infection can be inhibited by peptides derived from the heptad repeat regions of the human Parainfluenza Virus 3 [65]. This occurs as a result of the inhibition of 6 helix bundle formation, due to interactions between the native heptad repeat regions of NiV/Hev and peptide heptad repeats derived from Parainfluenza virus 3. The interaction of the heptad repeats of the human Parainfluenza Virus 3 with those of NiV/HeV along with their sequence conservation (Supporting Figure 1) could be suggestive of similarities in the post-fusion structure of these viruses, supporting our choice of template for modeling the post-fusion conformation of the F protein.

#### Modeling the M protein dimer

The M protein in NiV is crucial in initiating the budding of the virus. This protein homodimerizes before homo-oligomerizing and forming the viral matrix [66]. The crystal structure (PDB id: 4G1G) of a dimer of another Paramyxovirus, the Newcastle Disease Virus was used as a template for building a homology model of the M protein dimer. The target-template sequence identity was 19%, going up to 27% at the interface (29 identical residues out of 70). The model was energy minimized with GROMACS using the Amber99SB-ILDN force field [28] and evaluated using our empirical knowledge based scoring scheme, PIZSA [67]. The dimer had a PIZSA Z score of 1.69, well above the binding threshold of 1.50.

We also attempted to build several host-pathogen protein complexes but none of the models were evaluated favorably by FoldX [68] (Supporting Section 1).

### 2. Design and stability of protein peptide inhibitor complexes

#### Peptide inhibitor of the post fusion F protein

Protein F contains two helical domains identified as HRA and HRB. The HRA domain forms coiled-coil trimer that associates with three helices of the HRB domain to form a 6-helix coiled-coil bundle (sometimes referred to as 6HB) [69] (Figure 1), which is essential for its fusion with the host membrane. One strategy to inhibit the formation of this 6HB hexamer (which in turn would prevent the fusion of the host and viral membranes), is to design a peptide that would competitively bind to HRA domains, preventing its binding to the HRB helices. The 6HB forming regions of HRA and HRB, have heptad repeat sequence pattern [70] of a-b-c-d-e-f-g, such that hydrophobic amino acids occupy positions a/d and charged amino acids occupy e/g positions (Figure 1-A, B). An amino acid sequence of the inhibitor (IKKSKSYISKAQELL) was designed to mimic the HRB domain (LQQSKDYIKEAQRLL) such that all hydrophobic amino acids occupy a/d heptad positions (Figure 1-C, D). Further, the inhibitor sequence was designed to ensure that the atomic density in the core was optimized, similar to that observed in other coiled-coil proteins (unpublished study). Effectively, this meant changing the N terminal Leu in HRB to Ile in the inhibitor. Other amino acid replacements were done to ensure salt bridging between the inhibitor and HRA domain (Figure 1-D). Amino acids at non a/d heptad positions of the inhibitor that are not involved in interactions with the HRA domains were replaced by Lys. This is to introduce interactions between these Lys residues of the inhibitor with the Glu residues of the HRA domain (Figure 1-D). All other positions without any interacting partner on the HRA domain were replaced by Ser, to increase solvent interactions. The thirteenth residue of the inhibitor was changed from Arg to Glu to increase interactions with Lys on the HRA domain. The heptad repeat guided alignment of the inhibitor and 6HB domain of the HRB was used to structurally model the inhibitor using MODELLER v9.17.

**Figure 1.**
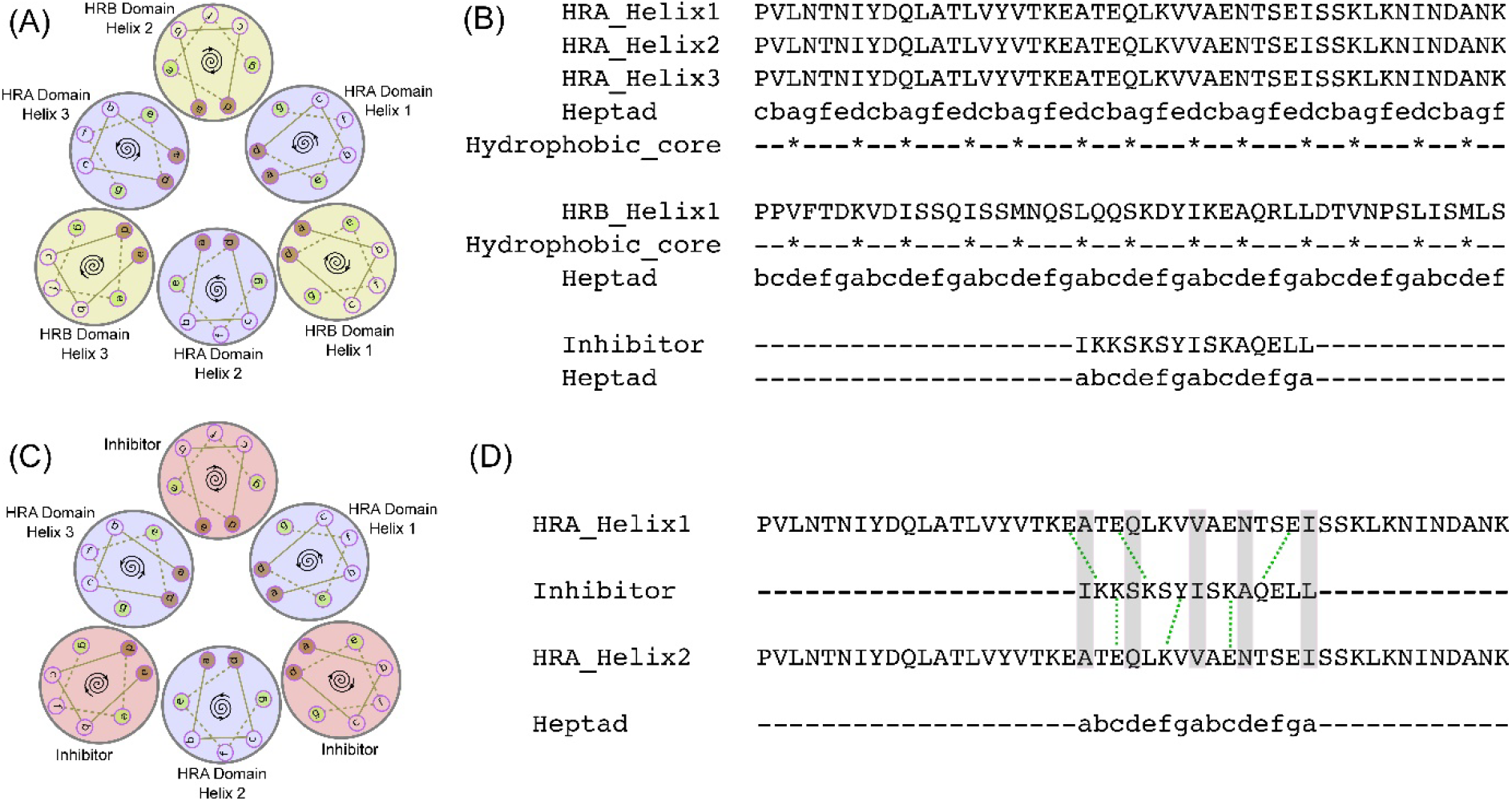
A) Heptad repeat representation of the 6-HB domain formed by HRA (purple circles) and HRB domain (yellow circles). Helices are represented as circles with an anticlockwise or clockwise spiral at the center showing the handedness of the helix. Amino acid heptad repeat positions are labeled with letters a through to g with hydrophobic amino acids occupying a and d positions. B) Heptad repeat assignment of HRA and HRB domain helices along with the designed inhibitor. Inhibitor heptad positions were assigned identical to the HRB domain. C) Heptad repeat representation of HRA (purple circles) domain and bound inhibitor (pink circles) replacing HRB domain. D) Salt bridges (green dotted lines) and interactions (gray bar) between the residues of the inhibitor and the HRA domain.

#### Peptide inhibitor of the M protein dimer

The binding sites on a monomer of M protein were detected with DEPTH [43] using default parameters. The predicted binding site that overlapped with the interface of the M protein dimer was used to target the dimerization process. The residues (RRTAGSTEK) of one monomer that interact with the predicted binding site of the other monomer at the dimer interface were modified by manual intervention. The last two amino acids (Glu-Lys) of the dimer interface sequence were modified to Ile-Asn such that they make specific interactions with the M protein. A 2 ns simulation with the unmodified sequence showed high fluctuations due to bulky charged groups at the C terminus. The C terminal Lys of the peptide is in close proximity to an Arg 197 on the M protein that leads to charge repulsion causing instability of the unmodified construct. Hence Lys was modified to Asn to reduce the size and repulsive forces. The penultimate residue, Glu was modified to Ile to improve hydrophobic contacts with its neighbors on the M protein. The modified peptide RRTAGSTIN was used as a putative M protein dimerization inhibitor for further analysis. Prevention of M protein dimerization could potentially prevent the virus from budding out of cells.

#### Peptide Inhibitors of G protein-ephrin interaction

The NiV infection is initiated by the binding of the G protein to the ephrin receptors on the host cell [71] (PDB ID: 2VSM). Inhibiting this protein-protein interaction could prevent viral entry. In this study, we have tested the feasibility of using 2 peptides to inhibit the G-protein – ephrin interaction. One peptide (FSPNLW) is the part of the ephrin-B2 receptor that interacts with the G-protein [72]. The other peptide (LAPHPSQ) is a part of a monocolonal antibody, m102.3, that binds [73] to both NiV and Hendra virus. A crystal structure of the antibody bound to Hendra virus G protein (PDB ID: 6CMG) was used as a template (79% target-template sequence identity) to construct the antibody-NiV G protein complex. 3D structural models of the speculated G-protein—peptide interactions were also constructed using MODELLER v9.17.

#### Computational prediction of the stability of the protein-inhibitor complexes

Three independent MD simulations of 100 ns each were performed to assess the stability of each of the four protein-peptide complexes. The peptide inhibitors designed against F and M proteins bind a hydrophilic pocket while the binding interactions of the G protein to its inhibitor are predominantly hydrophobic. For each of the trajectories, the total potential energy, the distance between the center of the protein and peptide, RMSD and RMSF of the peptide after superimposition of protein were analyzed and found to be consistent across independent runs (Supporting figure 2-5 and table 2-7). The F- and M-peptide complexes are stabilized by hydrogen bonds. A few of them (3 and 2 hydrogen bonds in F and M complexes respectively) (Supporting Table 3 and 5) are retained on average in over 50% of the trajectories. Hydrogen bond analysis was not done for the G protein - peptide inhibitor complexes since their binding is mediated mainly by hydrophobic interactions and there were no stable hydrogen bonds. The protein-peptide complex was stable during the simulation as can be inferred by the peptide RMSDs, peptide RMSFs and the distances between the protein and peptide. The distance of the center of the protein to that of the peptide fluctuated with a standard deviation of 0.03-0.09 nm (Supporting Tables 2, 4, 6, 7 and Supporting Figures 2, 3, 4, 5) around the average distance. For an explanation on the variations to these general observations, refer to Supporting Section 2. While these measures are all indicative of tight binding, we used the trajectories to determine the binding energy of association using the MM/PBSA protocol. The inhibitors of the F and M proteins bind tightly (*~*110 kJ/mol) to their targets (Supporting Tables 2, 4, 6, 7). However, in case of G protein inhibitors, the inhibitors FSPNLW and LAPHPSQ bind the G protein with *~*100 and ~60 kJ/mol, respectively, suggesting that ephrin-B2 receptor based design binds 40 kJ/mol stronger. This trend is also reflected in RMSD/RMSF values (Supporting Figures 4 and 5).

### 3. Prediction of putative small molecules that can bind to NiV proteins

We used 5 proteins (G, N, P, F and M) with experimental structures or models with high sequence coverage (~90%) in docking studies to predict plausible small molecule inhibitors (drugs or drug like molecules) against them. First, we predicted the plausible binding pockets on each of the proteins using the DEPTH server. A total of 12 binding pockets were predicted in G (2), N (4), P (2), F (1) and M (3) proteins. (Supporting Table 8). Two of the predicted binding pockets, one on the M protein and another on the G protein, are on the dimer interface and host protein (ephrin receptor) binding interface respectively. As mentioned in Section 2, these sites are important drug targets. All 12 binding sites were used to screen 22685 drug like molecules from the 70% redundant ZINC database of clean drug like molecules using two different docking tools, DOCK6.8 and Autodock4. The docking tools provide a docking energy score that was used to select possible high affinity binders. In the absence of an objective measure or threshold to determine strong binders, we chose the top 100 best scoring ligands for each of the pockets from both the docking tools. We then compared the two lists for common molecules. 70 molecules are identified by both Dock6.8 and Autodock4 for G (1), N (40), P (30), F (6) and M (37) proteins (Supporting Table 9). The grid score for the predicted complexes range between −71 to −32 units for DOCK6.8. The binding free energy for the predicted complexes range between −14 kcal/mol to −6 kcal/mol for Autodock4 (Supporting Table 9).

To corroborate our predictions, we measured the RMSD between the same ligand (in the common list) as docked by the two different tools (top 5 poses predicted by Autodock4 were compared to the top pose predicted by DOCK6.8), after superimposing the proteins. This measure is referred to as RMSD_lig. 10 drug-like molecules in N(5), P(4) and M(1) had an RMSD_lig less than 0.15 nm between their docked poses. In addition to conformational similarity, we also assessed the similarities in ligand-protein interactions, primarily hydrogen bonding (Supporting Table 10). Further, the hydrogen bonding interactions were ~50 % conserved in 5 of these complexes (with RMSD_lig < 0.15 nm). In a few instances, though the hydrogen bonding was not precisely the same, visual inspection of the complexes suggest that these bonds could be formed with small conformational changes. In 17 of the 70 cases the ligand was bound to the same pocket (RMSD_lig less than 0.2 nm).

Interestingly, a known drug (ZINC04829362), an antiasthmatic and antipsoriatic among other uses [74], binds to a pocket of the N protein with RMSD_lig of 0.085 nm. Another drug (ZINC12362922) used in the treatment of depression and Parkinson’s disease [75] also binds the N protein with RMSD_lig < 0.15 nm. The molecule with the best RMSD_lig (0.074 nm) from our screening, ZINC94258558 (Figure 2-A), also binds the N protein (Supporting Table 9).

**Figure 2.**
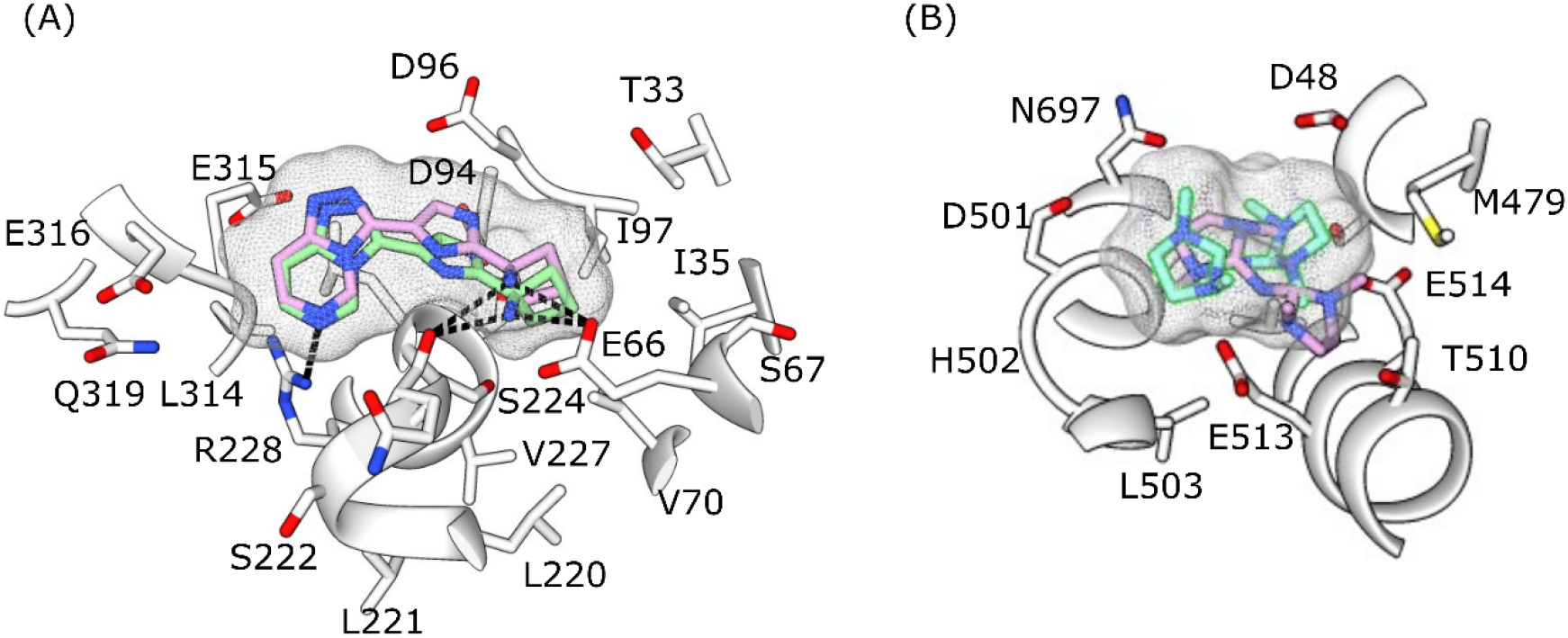
The docked poses of ZINC94258558 bound to N protein (A) and ZINC91252717 bound to P protein (B) as predicted by Autodock (green sticks with surface mesh) and Dock (lilac sticks with surface mesh). The protein is represented in white ribbons with the residues interacting with ligand shown in stick representation. Hydrogen bonds (only displayed in A) are shown as dashed lines.

After docking, we have shorted listed only those molecules whose RMSD_lig < 0.15 nm. There are however a few molecules that are of interest despite their relative large RMSD_lig values. The molecule ZINC91252717 is predicted as the best binder to P protein by Autodock4 (binding energy of −14 kcal/mol) and the second best binder by DOCK6.8 (grid score of −71) (Figure 2-B). These scores were among the best achieved during this docking exercise. Another molecule (ZINC00814199) was docked onto the M protein and formed 14 and 8 hydrogen bonds with Autodock and Dock respectively. It was also within the top 14 ranked compounds by both methods. By visual inspection, it is apparent that with small conformational changes, the Dock pose could also get 14 hydrogen bonding interactions. Lastly, the hydrophobic molecule ZINC63411510 is predicted to bind the G protein on its ephrin-B2 binding interface. Though both docking methods identify this site, the docking poses are different (RMSD_lig of 0.8 nm). We hypothesize that the hydrophobic nature of the binding pocket and its size is contributing to the difference in docked poses. There are also a few instances of the same drug binding different pockets, both within the same protein and on other proteins (Supporting Table 11).

### 4. Sequence variations in NiV isolates

At the time of modelling the NiV proteins, the sequence data from the 2018 outbreak was not available [49]. Hence, all the modelling was done by considering that sequence of the Malaysian strain. We rationalized that as the Malaysian and Bangladeshi/Indian strains shared a high degree (79-99 %) of sequence similarity, structural models using sequences of one strain would be applicable to the other, which is the basis of comparative modeling. However, we wanted to assess whether the efficacy of the designed/proposed therapeutic molecules would be affected by observed sequence variations between the different strains (7 Malaysian, 3 Bangladeshi and 5 Indian) of NiV.

The amino acid variations (Supporting Table 12) were mapped onto their respective structures. All protein sequences are of equal length except the V protein whose lengths vary between the different strains. The V and W protein have the least sequence conservation (~79%) while the M protein is the most conserved (98.6%). A general observation is that the Bangladeshi and Indian strains are more similar to one another than they are to the Malaysian sequences (Supporting Figure 6).

We mapped the sequence variations onto all the protein structures/models that were used for peptide inhibitor design and drug docking. No variations in the sequence were found close to the peptide inhibitor binding sites on the F, M and G proteins. We found 1 (Lys236Arg), 2 (Asp188Glu, Gln211Arg), 1 (Asp252Gly) and 1 (Ile331Val) variations close to the docking sites on G, N, F and M protein respectively. ZINC63411510, ZINC63411510 (bound to G protein), ZINC20163996 (bound to F protein) and ZINC72131030 (bound to M protein) and all the small molecules bound to PN4 binding site of N were within 0.5 nm of the mutated amino acids. All the mutations (except for Asp252Gly on F protein) on the binding site were conservative (similar physico-chemical properties and BLOSUM62 score >= 0) and hence are conjectured not to affect the interactions between the protein and the inhibitor. Though there is a non-conservative change (ASP252Gly) in one of the drug/inhibitor binding sites of the F protein, this position is not involved in H-bonding with the ligand. Hence the binding of the inhibitor to the protein is probably not going to be affected. No single sequence variant we have studied appears to show that the drug binding would be directly affected.

### 5. Web service and Database

We have archived all structures/models of NiV proteins and their inhibitor bound complexes in a consolidated database at http://cospi.iiserpune.ac.in/Nipah. The data at this site lists details of modeling, docking features and multiple sequence alignment (between the various NiV strains) such as template PDB code, target-template sequence identity, model quality assessment score, docking energies, docking rank and the RMSD_lig between the docking poses

## Discussions

NiV is a deadly zoonotic virus with a mortality rate of 72% and 86% in Bangladesh and India respectively. There is no known drugs/therapeutics against NiV. The overarching aim of this study is to computationally design inhibitors and predict small molecule drugs against NiV proteins. To design/predict therapeutic molecules to act against NiV, we characterized all of its proteins. As a part of this effort, we constructed partial models of 5 NiV proteins *viz*., M, L, V, W proteins along with the post fusion conformation of the F protein. The structure of the post-fusion conformation of the F protein is modeled for the first time in this study. Our model is based on the post-fusion structures of another class I fusion protein from Human Parainfluenza virus 3.

Our efforts have increased the coverage of existing structures of the G, N and P proteins (by 13%, 8% and 8% respectively) by modeling a fraction of their unresolved residues. No reliable models could be generated for the C protein. Effectively, we doubled the number of amino acids in the NiV proteome that were structurally characterized. While our aim is to use these models to predict/design inhibitors, we believe that many of our models are by themselves quite insightful. They could serve as templates for future structure-guided drug designing efforts against members of the Paramyxoviridae family. We attempted to build complexes of the viral and host protein (host cathepsin-L with NiV F protein and host AP3-B1 with NiV M protein) to target the interactions for inhibitor design. However, we were unsuccessful in making reliable models of host-pathogen protein-protein interaction complexes. With improvements to protein-protein docking methods, the quality of such models of complexes could be improved, which in turn would help in better targeting host-viral interactions.

We next used these models to design 4 peptide inhibitors against the F, M and G proteins. The inhibitor against F protein would putatively prevent the pre to post fusion transition of the F protein, a crucial step for viral entry. Our model of the post fusion conformation of the F protein was crucial in designing this inhibitor. Another inhibitor against the M protein was designed such that it would prevent the dimerization of the protein, hence preventing the budding process. The two inhibitors against the G proteins were selected such that they bind to the ephrin receptor binding pocket, preventing viral attachment to the host cell. The peptides here mimic the ephrin-B2 protein and an antibody (m102.3) that are bound at the same site. We conjectured that these peptides would competitively inhibit the G protein from binding the host ephrin receptors. All of these protein-peptide systems were subjected to triplicate runs of 100 ns MD simulations to assess interaction strengths. The distance of the center of the inhibitor and the peptide fluctuates with a standard deviation of 0.03-0.09 nm from the mean distance, indicative of the inhibitor remaining in the binding pocket. The inhibitors against the F and M proteins also had stable hydrogen bond associations in the MD trajectories. Binding affinity calculations suggest that three of the designed putative inhibitors bind tightly (*~*100 kJ/mol) to their targets, making them promising leads against NiV proteins.

We screened a set of drug like molecules in a docking exercise to identify potential small molecule inhibitors of NiV. The screen consisted of 22685 compounds of the 70% non-redundant set of clean drug like molecules of the ZINC library. The docking onto the NiV proteins was done using two different docking programs, Autodock4 and Dock6.8. Empirically, we chose the top 100 ligands from each of the two methods and selected those that were common between them. This resulted in 70 compounds that bound the G, N, P, F and M proteins of NiV. As a more stringent test, we whittled down this list to only include those molecules that were docked in similar poses (empirically chosen RMSD of 0.15 nm or smaller) on the same binding site. Hence, we predicted 10 compounds that would inhibit the N (5), P (4) and M (1) proteins of NiV. In addition we also included 3 drugs to the list that did not clear the criteria explained above. These drugs include one that binds the G protein on its ephrin binding interface and two others which bind to P and M proteins. The most important aspect of the docking study is that the molecular screen consists of known drugs or drug like compounds. The implication is that a few of our proposed inhibitors could be readily tested and repurposed. For instance, we have identified Cyclopent-1-ene-1,2-dicarboxylic acid (ZINC04829362) as an inhibitor of the NiV N protein. This compound is a known drug prescribed for antiasthmatic and antopsoriatic among other disorders. Another example is Bicyclo[2.2.1]hepta-2,5-diene-2,3-dicarboxylic acid (ZINC12362922) that we propose also inhibits the N protein, is a drug prescribed against depression and Parkinson’s disease. We cannot overemphasize the importance of these computational predictions, especially for swift acting potent viruses as NiV where mortality rates are high.

Finally, we assessed the how effective our proposed inhibitors would be against different strains of the virus and assess the risk of the virus getting drug resistant. For this, we studied 3 Bangladeshi, 7 Malaysian and 5 Indian strains and inferred the variations between the various strains from their multiple sequences alignment. Further, we investigated whether such changes would affect inhibitor binding. Here, we narrowed the changes only to those residues that were in direct contact (< 0.5 nm) from the inhibitors. We precluded the possibility of allosteric interactions. None of the residues contacting the peptide inhibitors showed any variations in their sequence. Only 5 residue positions that were involved in binding the drug like inhibitors were changed between the different strains. 4 of these changes are conservative substitutions where the nature of the mutated residue is not deemed to change the binding property of the protein to its inhibitor. Only 1 amino acid change of Asp252Gly of the F protein is a non-conservative change, however the Asp is not involved in hydrogen bonding with the ligand. We conclude that it is likely that the proposed inhibitors would be potent against all strains of the virus

Nipah and other zoonotic virus pose a serious epidemic threat. Computational approaches can help identify/design inhibitors that could be rapidly tested or even deployed as they may be drugs previously licensed for other uses. Our study also has connotations for related viruses such as Hendra and other Paramyxoviruses. Importantly, our models and the web pages we have created could be modified to serve as portal to study the epidemiology of the virus should there be further outbreaks.

## Acknowledgements

The authors would like to acknowledge Swastik Mishra, Yogendra Ramtirtha and Kundan Kumar for useful discussions.

## Supporting Information

Supporting table 1- Model quality evaluation of the protein structures built using I-TASSER web server. The best model predicted by I-TASSER (based on their C-Score) have their Normalized DOPE scores and C-scores in bold. TM-scores and RMSDs are only calculated for the best models. L protein was divided into three domains, indicated by their residue numbers in parentheses, and modeled separately.

Supporting Table 2- Mean and standard deviation of the energy, distance of the center of the inhibitor with the center of the F protein, number of hydrogen bonds between the inhibitor and the protein, RMSD of the inhibitor and the protein-peptide binding energies obtained from the three 100ns MD simulations of F protein-inhibitor complex.

Supporting Table 3- Percentage of the snapshots with hydrogen bonds between the chain D of inhibitor with chain C and E of the F protein.

Supporting Table 4- Mean and standard deviation of the energy, distance of the center of the inhibitor with the center of the M protein, number of hydrogen bonds between the inhibitor and the M protein, RMSD of the inhibitor and the protein-peptide binding energies obtained from the three 100ns MD simulations of the M protein-inhibitor complex.

Supporting Table 5- Percentage of snapshots with hydrogen bonds between chain B of the inhibitor and chain A of the M protein.

Supporting Table 6- Mean and standard deviation of the energy, distance of the center of the FSPNLW inhibitor with the center of the G protein, RMSD of the inhibitor and the protein-peptide binding energies obtained from the three 100 ns MD simulations of G protein-FSPNLW inhibitor complex.

Supporting Table 7- Mean and standard deviation of the energy, distance of the center of the LAPHPSQ inhibitor with the center of the G protein, RMSD of the inhibitor and the protein-peptide binding energies obtained from the three 100 ns MD simulations of G protein-LAPHPSQ inhibitor complex.

Supporting Table 8- List of pocket lining residues for each pocket of NiV Proteins. The residue name is followed by the residue number. The chain id has been depicted after the dot.

Supporting Table 9- List of the ranks and energy values of the small drug like molecules that were predicted in the top 100 by both DOCK6.8 and Autodock4. RMSD_1 – RMSD_5 are the RMSDs of the 5 best Autodock4 poses with the best scoring Dock6.8 pose. The least RMSD is depicted in bold. Cells highlighted in yellow have RMSDs better than 0.15 nm. Pocket number indicates pockets from Autodock. Some of the Autodock pockets have been subdivided by DOCK, which indicates the subsections in each pocket.

Supporting Table 10- Number of Hydrogen bonds that are formed between the selected pose for DOCK6.8 and Autodock4 with the protein. Number of common Hydrogen bonds indicates the number of hydrogen bonds that are common between the predicted poses of the ligand from Autodock4 and DOCK6.8.

Supporting Table 11 – Same drug like molecule predicted to bind different pockets of the same or different protein. The binding pocket has been mentioned in parenthesis.

Supporting Table 12 – The sequence variations between the 15 NiV strains. The mutations are mentioned by the residue number followed by the amino acids present in different strains.

Supporting figure 1 - Conformational change of the human Parainfluenza Virus 3 (HPIV3) fusion protein and its sequence conservation with Nipah Virus (NiV) and Hendra Virus (HeV). The fusion protein undergoes a large conformational change from the pre-fusion state (A, PDB id: 6MJZ) to post-fusion state (B, PDB id: 1ZTM) to form the 6 helix bundle by interactions between the HRA domain (Salmon ribbon) and HRB domain (Cyan ribbon) heptad repeat regions. (C) Alignment of the heptad repeat regions between fusion protein sequences of the three viruses (Uniprot ids - HPIV3: P06828, NiV: Q9IH63, HeV: O89342). The alignment is color coded based on ClustalX.

Supporting Figure 2- A) Energy of the F protein-inhibitor complex during 100 ns of MD simulation B) Distance of the center of the inhibitor from the center of the F protein during the simulation C) Number of hydrogen bonds between the F protein-inhibitor complex during the simulations D) Plot showing the formation of hydrogen bonds between inhibitor and F protein over 100 ns trajectories. Y axis shows the 11 different hydrogen bonds identified as numbered index (Supporting Table 3). X axis labels time instant during simulation. Each rectangular color box represent presence of hydrogen bond for a particular run. E) Root mean square deviation (RMSD) ^#^ of the designed inhibitor during the simulations F) Root mean square fluctuation (RMSF) ^#^ of the inhibitory peptide during the simulations. Each of the simulations were run in triplicate, with each run being color coded as red, green and blue.

Supporting Figure 3- A) Energy of the M protein-inhibitor complex during 100 ns of MD simulation B) Distance of the center of the inhibitor from the center of the M protein during the simulation C) Number of hydrogen bonds between the M protein-inhibitor complex during the simulations D) Plot showing the formation of hydrogen bonds between inhibitor and M protein over 100 ns trajectories. Y axis shows the 8 different hydrogen bonds identified as numbered index (Supporting table 5). X axis labels time instant during simulation. Each rectangular color box represents presence of hydrogen bond for a particular run. E) RMSD ^#^ of the designed inhibitor during the simulations F) RMSF ^#^ of the inhibitory peptide during the simulations. Each of the simulations were run in triplicate, with each run being color coded as red, green and blue.

Supporting Figure 4- A) Energy of the G protein-FSPNLW inhibitor complex during 100 ns of MD simulation B) Distance of the center of the inhibitor from the center of the G protein during the simulation C) RMSD ^#^ of the designed inhibitor during the simulation D) RMSF ^#^ of the inhibitory peptide during the simulation. Each of the simulation were run in triplicate, each run being color coded as red, green and blue.

Supporting Figure 5- A) Energy of the G protein-LAPHPSQ inhibitor complex during 100 ns of MD simulation B) Distance of the center of the inhibitor from the center of the G protein during the simulation * C) RMSD ^#^ of the designed inhibitor during the simulation D) RMSF ^#^ of the inhibitory peptide during the simulation. Each of the simulation were run in triplicate, each run being color coded as red, green and blue.

Supporting Figure 6 – Heatmap showing the sequence conservation between the different strains of NiV for (A) C protein (B) F protein (C) G protein (D) L protein (E) M protein (F) N protein (G) P protein (H) V protein (I) W protein. The color gradient represents sequence conservation where white indicates 100% conservation and redder shades indicate lesser sequence conservation. The labelling convention is Protein_Country_Genome-accession code.

Supporting Section 1 – Modeling of host-pathogen interactions

Supporting Section 2 – Molecular dynamics simulations of protein-peptide inhibitor complexes

