## Supplementary Data for "Predicting and Designing therapeutics against the Nipah virus"

Supporting table 1- Model quality evaluation of the protein structures built using I-TASSER web server. The best model predicted by I-TASSER (based on their C-Score) have their Normalized DOPE scores and C-scores in bold. TM-scores and RMSDs are only calculated for the best models. L protein was divided into three domains, indicated by their residue numbers in parentheses, and modeled separately.

| **Protein** | **Normalized DOPE** | **C-score** | **Predicted TM-score^$^** | **Predicted RMSD** |
| --- | --- | --- | --- | --- |
| V | **2.09**, 1.70, 1.99, 1.23, 1.27 | **-0.79**, -1.82, -0.32, -3.56, -2.69 | 0.61 | 8.9 |
| W | **1.45**, 0.77, 0.75, 1.56, 0.80 | **-1.42**, -1.73, -3.07, -4.34, -3.30 | 0.54 | 10.4 |
| C | **0.49**, -0.08, -1.53, -0.33, -0.88 | **-3.68**, -3.67, -3.29, -4.16, -4.09 | 0.31 | 13.6 |
| L^#^ (14 - 1177) | **0.29**, 0.31, -0.04, 0.01, -0.16 | **0.07**, -0.30, -1.87, -1.05, -0.83 | 0.72 | 9.1 |
| L^#^ (1191 - 1435) | **0.52** | **1.1** | 0.86 | 3.6 |
| L^#^ (1553 - 1859) | **0.21**, 0.82, 0.95, 2.69, 0.44 | **-2.61**, -4.24, -4.52, -4.76, -5.00 | 0.41 | 12.4 |

^#^The protein was built domain wise because I-TASSER has a maximum size limit of 1500 residues.

^$^Although models built for V, W proteins and two of the Polymerase L domains had a TM-scores greater than 0.5, none of these models had a Normalized DOPE score less than or equal to zero and therefore were not used further in the study.

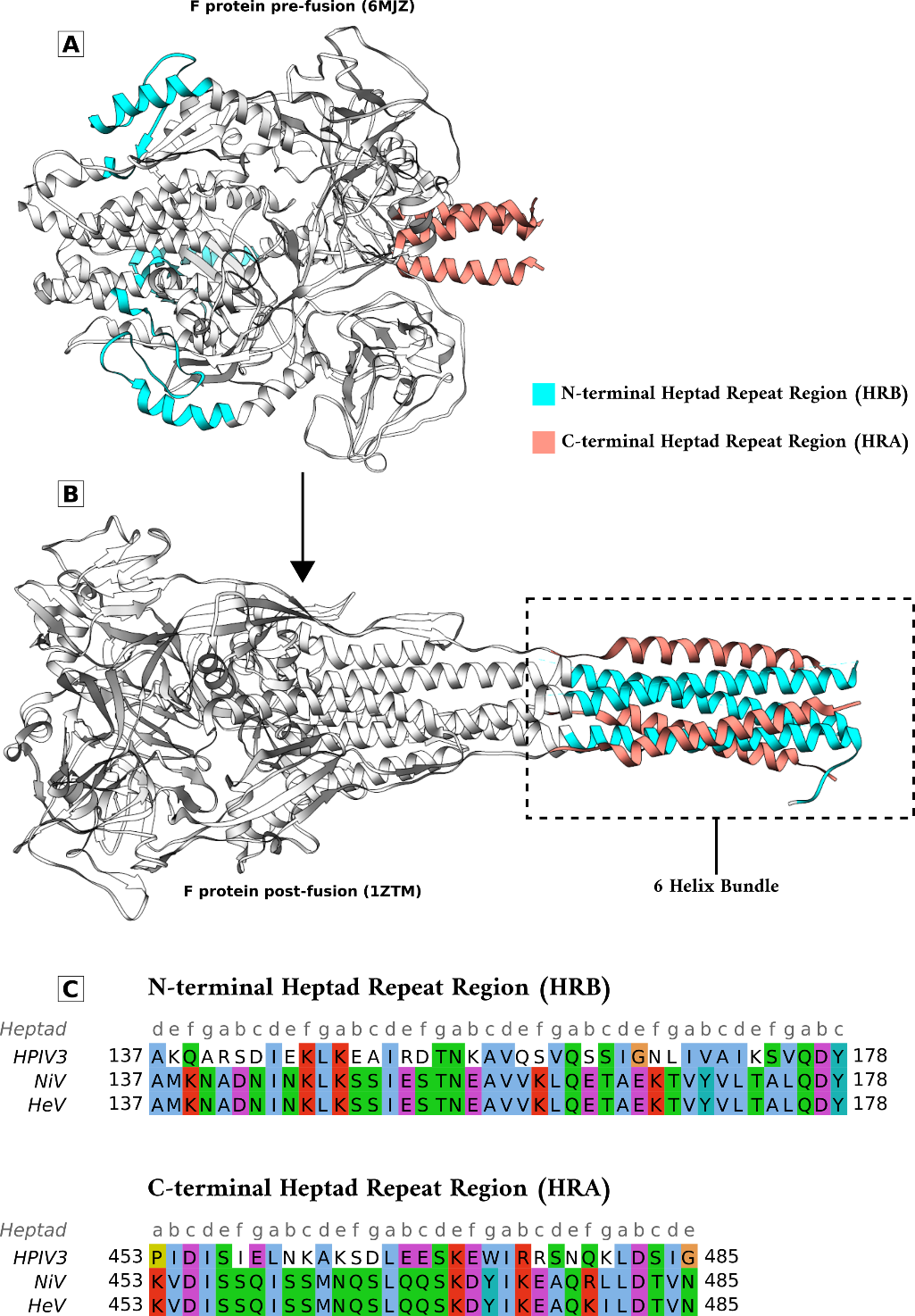

Supporting figure 1 - Conformational change of the human Parainfluenza Virus 3 (HPIV3) fusion protein and its sequence conservation with Nipah Virus (NiV) and Hendra Virus (HeV). The fusion protein undergoes a large conformational change from the pre-fusion state (A, PDB id: 6MJZ) to post-fusion state (B, PDB id: 1ZTM) to form the 6 helix bundle by interactions between the HRA domain (Salmon ribbon) and HRB domain (Cyan ribbon) heptad repeat regions. (C) Alignment of the heptad repeat regions between fusion protein sequences of the three viruses (Uniprot ids - HPIV3: P06828, NiV: Q9IH63, HeV: O89342). The alignment is color coded based on ClustalX.

**Supporting Section 1- Modeling of host-pathogen interactions**

We attempted to model 2 host-pathogen protein complexes involving human cathepsin L with viral F protein and human AP3-B1 with viral M protein. The interacting interfaces of the host-viral complexes are potential targets for designed therapeutics. Human cathepsin-L interaction with F protein is crucial for activation of the F protein to initiate fusion [1]. Viral assembly requires the interaction of M protein with host AP3-B1 [2]. HADDOCK [3], Patchdock [4,5] and Galaxy [6] docking servers were used to model these protein-protein complexes. However, the resulting models had a positive free energy of binding as calculated with FoldX [7], and were therefore not used further in the study.

**Section 2 – Molecular dynamics simulation of protein-peptide inhibitor complexes**

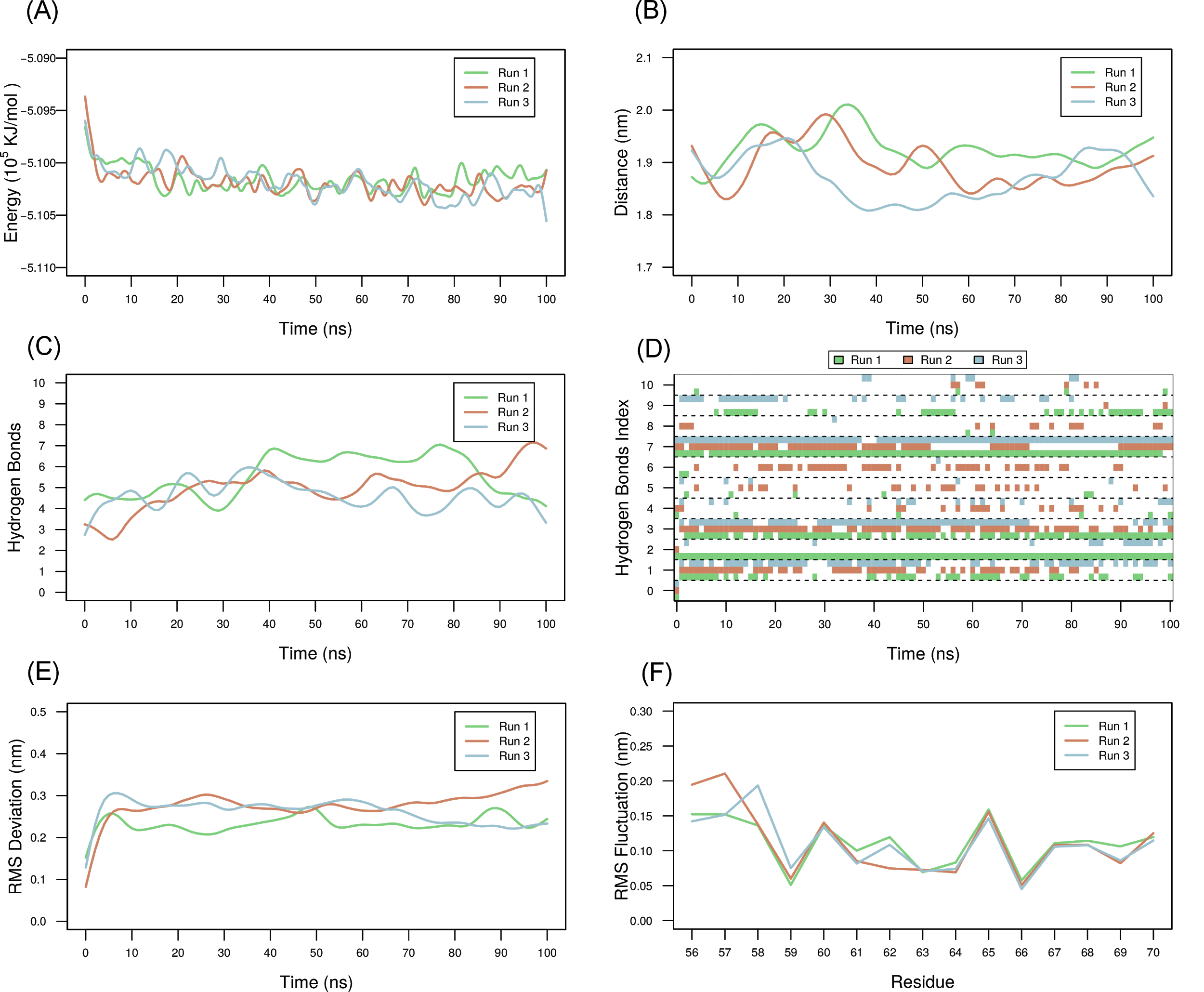

Supporting Figure 2- A) Energy of the F protein-inhibitor complex during 100 ns of MD simulation B) Distance of the center of the inhibitor from the center of the F protein during the simulation C) Number of hydrogen bonds between the F protein-inhibitor complex during the simulations D) Plot showing the formation of hydrogen bonds between inhibitor and F protein over 100 ns trajectories. Y axis shows the 11 different hydrogen bonds identified as numbered index (Supporting Table 3). X axis labels time instant during simulation. Each rectangular color box represent presence of hydrogen bond for a particular run. E) Root mean square deviation (RMSD) ^#^ of the designed inhibitor during the simulations F) Root mean square fluctuation (RMSF) ^#^ of the inhibitory peptide during the simulations. Each of the simulations were run in triplicate, with each run being color coded as red, green and blue.

^#^ RMSD and RMSF were calculated for the inhibitor by superimposing the protein molecule

Supporting Table 2- Mean and standard deviation of the energy, distance of the center of the inhibitor with the center of the F protein, number of hydrogen bonds between the inhibitor and the protein, RMSD of the inhibitor and the protein-peptide binding energies obtained from the three 100ns MD simulations of F protein-inhibitor complex.

| **Run** | **Energy (kJ/mol)** | | **Protein-peptide distance (nm)** | | **Number of Hydrogen Bonds** | | **RMSD (nm)** | | **Binding energies (kJ/mol)** | |
| --- | --- | --- | --- | --- | --- | --- | --- | --- | --- | --- |
|  | **Mean** | **SD** | **Mean** | **SD** | **Mean** | **SD** | **Mean** | **SD** | **Mean** | **SD** |
| 1 | -510162 | 1093 | 1.93 | 0.05 | 5.54 | 1.48 | 0.23 | 0.04 | -102.5 | 9.6 |
| 2 | -510194 | 1084 | 1.89 | 0.05 | 4.99 | 1.74 | 0.28 | 0.04 | -117.8 | 9.2 |
| 3 | -510179 | 1090 | 1.87 | 0.06 | 4.62 | 1.32 | 0.26 | 0.04 | -102.9 | 9.7 |
| **Mean** | **-510178** |  | **1.90** |  | **5.05** |  | **0.26** |  | **-107.7** |  |

Supporting Table 3- Percentage of the snapshots with hydrogen bonds between the chain D of inhibitor with chain C and E of the F protein.

| **Index** | **Hydrogen Bond Partner** | **Run 1 (%)** | **Run 2 (%)** | **Run 3 (%)** |
| --- | --- | --- | --- | --- |
| 0 | 56ILE.D-26THR.E | 1 | 1 | 1 |
| 1 | 56ILE.D-28GLU.C | 41.6 | 55.4 | 59.4 |
| 2 | 59SER.D-19ALA.E | 100 | 1 | 12.9 |
| 3 | 59SER.D-24GLN.C | 77.2 | 75.2 | 68.3 |
| 4 | 60LYS.D-24GLN.C | 5.9 | 18.8 | 17.8 |
| 5 | 65LYS.D-15SER.E | 5.9 | 22.8 | 5 |
| 6 | 65LYS.D-18GLU.E | 2 | 42.6 | 1 |
| 7 | 66ALA.D-12SER.E | 97 | 64.4 | 97 |
| 8 | 67GLN.D-14GLU.C | 2 | 12.9 | 1 |
| 9 | 67GLN.D-17ASN.C | 36.6 | 2 | 31.7 |
| 10 | 69LEU.D-8LYS.E | 4 | 6.9 | 6.9 |

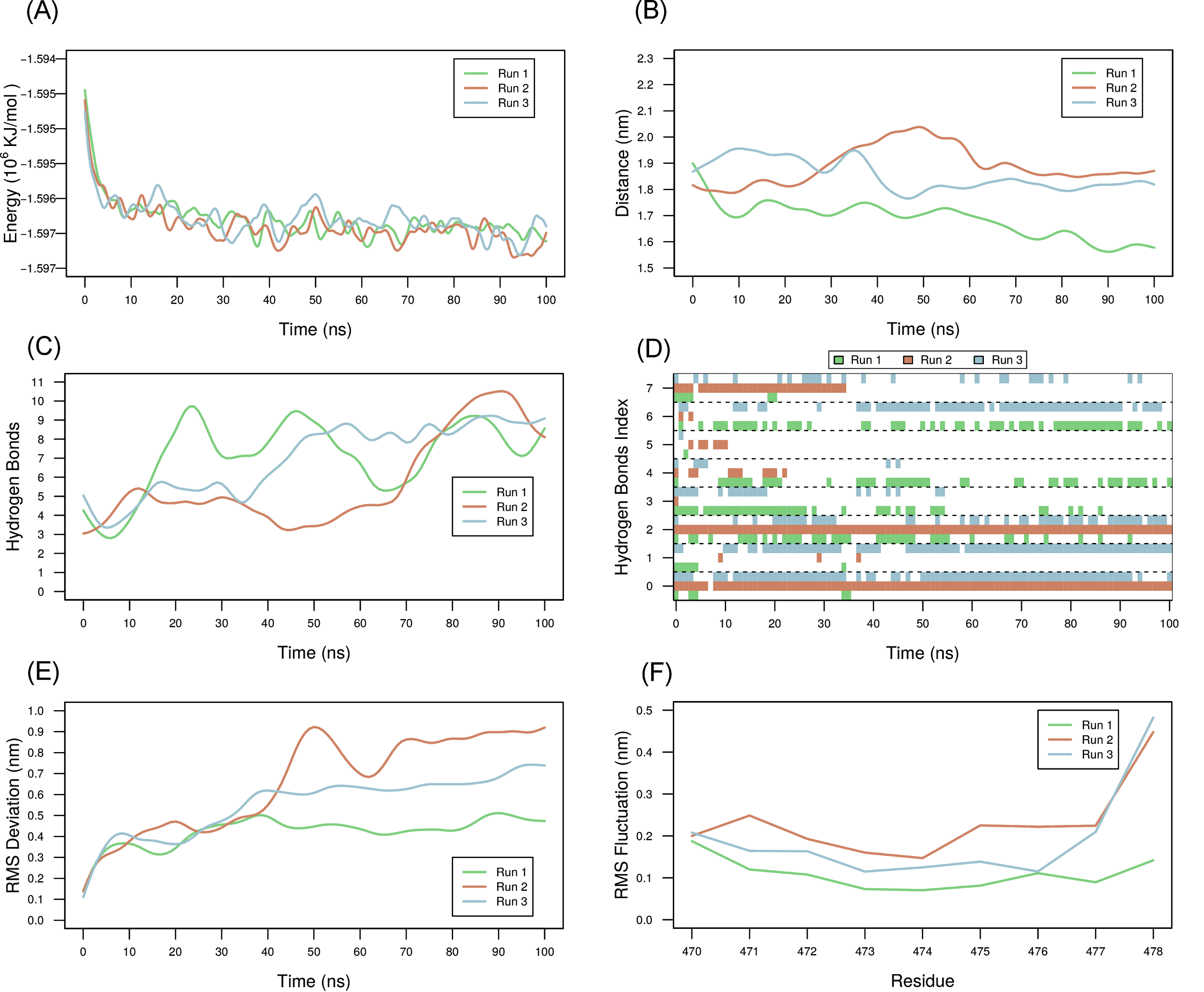

Supporting Figure 3- A) Energy of the M protein-inhibitor complex during 100 ns of MD simulation B) Distance of the center of the inhibitor from the center of the M protein during the simulation C) Number of hydrogen bonds between the M protein-inhibitor complex during the simulations D) Plot showing the formation of hydrogen bonds between inhibitor and M protein over 100 ns trajectories. Y axis shows the 8 different hydrogen bonds identified as numbered index (Supporting table 5). X axis labels time instant during simulation. Each rectangular color box represents presence of hydrogen bond for a particular run. E) RMSD ^#^ of the designed inhibitor during the simulations F) RMSF ^#^ of the inhibitory peptide during the simulations. Each of the simulations were run in triplicate, with each run being color coded as red, green and blue.

^#^ RMSD and RMSF were calculated for the inhibitor by superimposing the protein molecule

Supporting Table 4- Mean and standard deviation of the energy, distance of the center of the inhibitor with the center of the M protein, number of hydrogen bonds between the inhibitor and the M protein, RMSD of the inhibitor and the protein-peptide binding energies obtained from the three 100ns MD simulations of the M protein-inhibitor complex.

| **Run** | **Energy (kJ/mol)** | | **Protein-peptide distance (nm)** | | **Hydrogen Bonds** | | **RMSD (nm)** | | **Binding energies (kJ/mol)** | |
| --- | --- | --- | --- | --- | --- | --- | --- | --- | --- | --- |
|  | **Mean** | **SD** | **Mean** | **SD** | **Mean** | **SD** | **Mean** | **SD** | **Mean** | **SD** |
| 1 | -1596309 | 1851 | 1.69 | 0.08 | 7.15 | 2.41 | 0.42 | 0.08 | -107.9 | 11.6 |
| 2 | -1596399 | 1868 | 1.89 | 0.09 | 5.73 | 2.59 | 0.66 | 0.23 | -121.7 | 11.9 |
| 3 | -1596287 | 1856 | 1.85 | 0.07 | 6.98 | 2.29 | 0.56 | 0.14 | -98.4 | 12.0 |
| **Mean** | **-1596332** |  | **1.81** |  | **6.62** |  | **0.55** |  | **-107.7** |  |

Supporting Table 5- Percentage of snapshots with hydrogen bonds between chain B of the inhibitor and chain A of the M protein.

| **Index** | **Hydrogen Bond Partner** | **Run 1 (%)** | **Run 2 (%)** | **Run 3 (%)** |
| --- | --- | --- | --- | --- |
| 0 | 470ARG.B-335PHE.A | 5.0 | 99.0 | 80.2 |
| 1 | 470ARG.B-338TYR.A | 5.9 | 3.0 | 80.2 |
| 2 | 470ARG.B-340ASP.A | 40.6 | 100.0 | 45.5 |
| 3 | 471ARG.B-328GLN.A | 41.6 | 1.0 | 19.8 |
| 4 | 473ALA.B-197ARG.A | 44.6 | 9.9 | 5.9 |
| 5 | 475SER.B-194LEU.A | 1.0 | 5.9 | 1.0 |
| 6 | 475SER.B-195GLU.A | 53.5 | 2.0 | 62.4 |
| 7 | 476THR.B-304ASP.A | 5.9 | 33.7 | 24.8 |

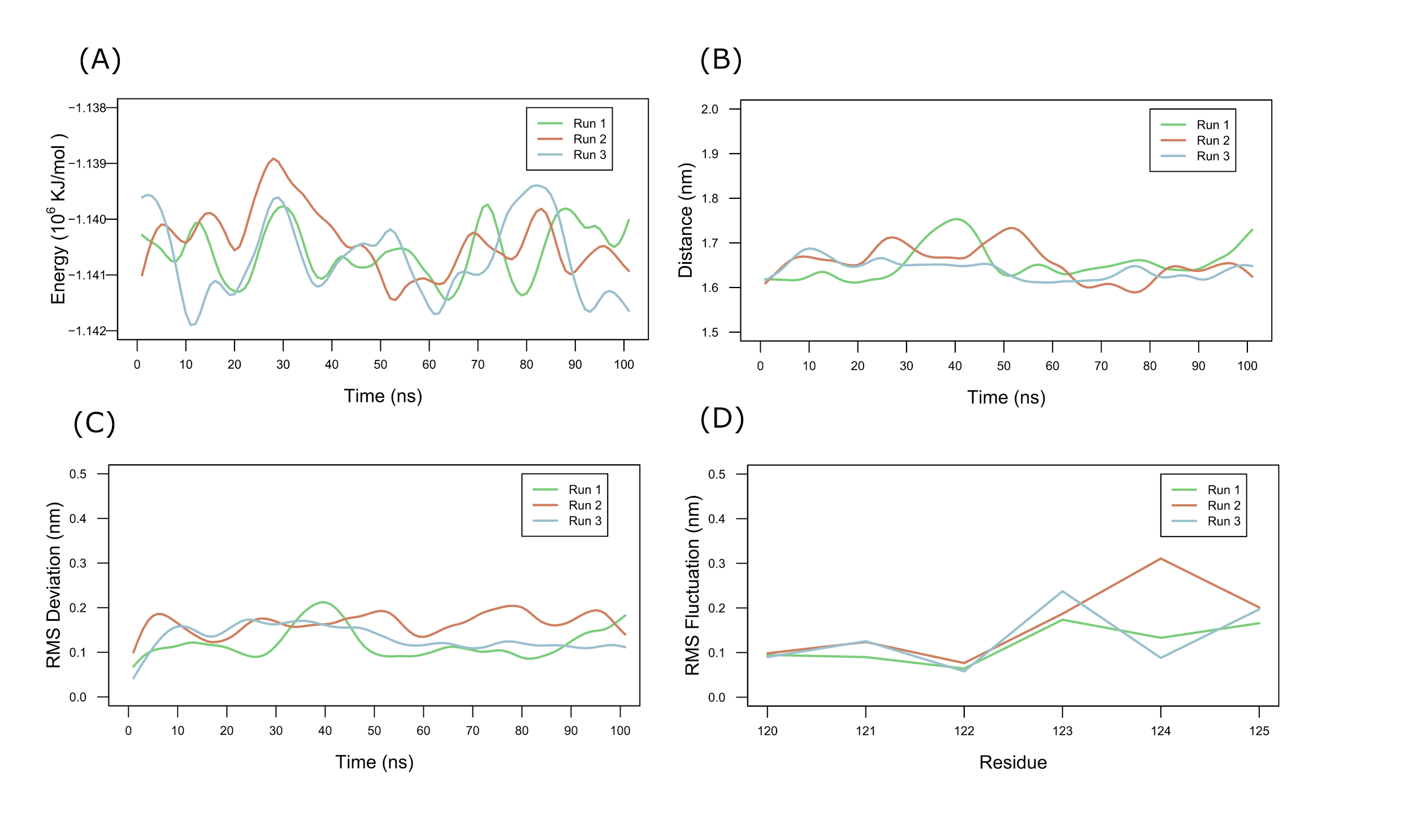

Supporting Figure 4- A) Energy of the G protein-FSPNLW inhibitor complex during 100 ns of MD simulation B) Distance of the center of the inhibitor from the center of the G protein during the simulation C) RMSD ^#^ of the designed inhibitor during the simulation D) RMSF ^#^ of the inhibitory peptide during the simulation. Each of the simulation were run in triplicate, each run being color coded as red, green and blue.

^#^ RMSD and RMSF were calculated for the inhibitor by superimposing the protein molecule

Supporting Table 6- Mean and standard deviation of the energy, distance of the center of the FSPNLW inhibitor with the center of the G protein, RMSD of the inhibitor and the protein-peptide binding energies obtained from the three 100 ns MD simulations of G protein-FSPNLW inhibitor complex.

| **Run** | **Energy (kJ/mol)** | | **Protein-peptide distance (nm)** | | **RMSD (nm)** | | **Binding energies (kJ/mol)** | |
| --- | --- | --- | --- | --- | --- | --- | --- | --- |
|  | **Mean** | **SD** | **Mean** | **SD** | **Mean** | **SD** | **Mean** | **SD** |
| 1 | -1140578 | 1550 | 1.65 | 0.05 | 0.12 | 0.04 | -107.2 | 10.8 |
| 2 | -1140385 | 1714 | 1.66 | 0.05 | 0.17 | 0.04 | -96.0 | 10.8 |
| 3 | -1140696 | 1708 | 1.64 | 0.03 | 0.13 | 0.03 | -93.9 | 12.2 |
| **Mean** | **-1140553** |  | **1.65** |  | **0.14** |  | **-99.0** |  |

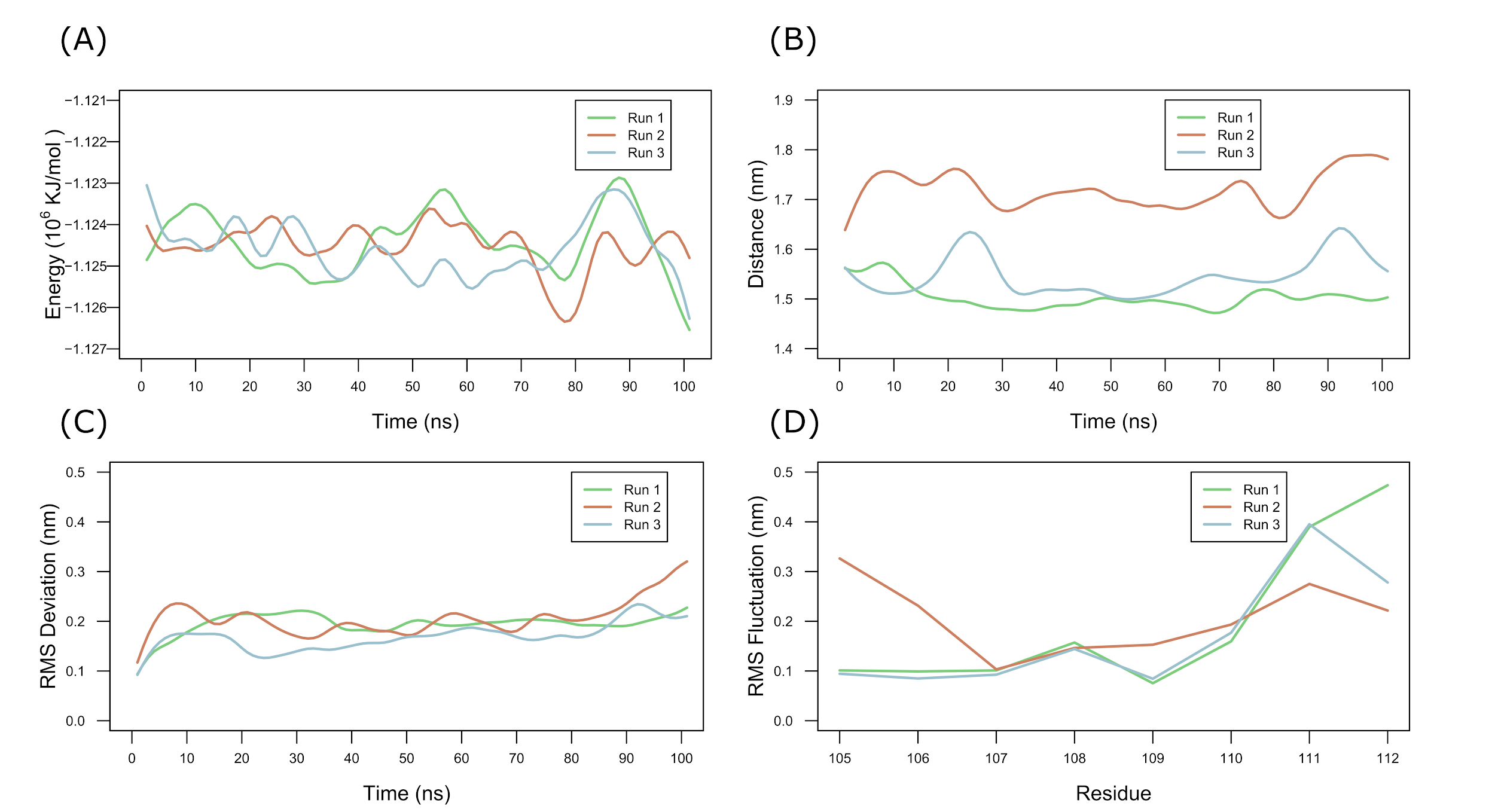

Supporting Figure 5- A) Energy of the G protein-LAPHPSQ inhibitor complex during 100 ns of MD simulation B) Distance of the center of the inhibitor from the center of the G protein during the simulation * C) RMSD ^#^ of the designed inhibitor during the simulation D) RMSF ^#^ of the inhibitory peptide during the simulation. Each of the simulation were run in triplicate, each run being color coded as red, green and blue.

* The distance between the inhibitor and the protein was consistently 0.2 nm higher for one of the replicates. This is because we considered the center of the protein as the all atom center and fluctuations in the side chains can explain the deviation.

^#^ RMSD and RMSF were calculated for the inhibitor by superimposing the protein molecule

Supporting Table 7- Mean and standard deviation of the energy, distance of the center of the LAPHPSQ inhibitor with the center of the G protein, RMSD of the inhibitor and the protein-peptide binding energies obtained from the three 100 ns MD simulations of G protein-LAPHPSQ inhibitor complex.

| **Run** | **Energy (kJ/mol)** | | **Protein-peptide distance (nm)** | | **RMSD (nm)** | | **Binding energies (kJ/mol)** | |
| --- | --- | --- | --- | --- | --- | --- | --- | --- |
|  | **Mean** | **SD** | **Mean** | **SD** | **Mean** | **SD** | **Mean** | **SD** |
| 1 | -1124416 | 1628 | 1.50 | 0.03 | 0.19 | 0.03 | -61.0 | 12.9 |
| 2 | -1124513 | 1695 | 1.72 | 0.05 | 0.21 | 0.05 | -58.9 | 11.4 |
| 3 | -1124582 | 1578 | 1.54 | 0.05 | 0.17 | 0.03 | -65.1 | 12.2 |
| **Mean** | **-1124504** |  | **1.59** |  | **0.19** |  | **-61.7** |  |

Supporting Table 8- List of pocket lining residues for each pocket of NiV Proteins. The residue name is followed by the residue number. The chain id has been depicted after the dot.

| **NiV protein** | **Pocket** | **Pocket lining residue numbers** |
| --- | --- | --- |
| Glycoprotein | PG1 | F458.A, W504.A, Q559.A, D219.A, Y280.A, L305.A, Q490.A |
|  | PG2 | P500.A, G489.A, R435.A, W479.A, S432.A, E430.A, R344.A, K376.A, F375.A, N378.A, S398.A, P383.A |
| Nucleoprotein | PN1 | S67.A, A65.A, V58.A, I131.A, L128.A, E124.A, R36.A, F38.A, K34.A |
|  | PN2 | K69.A, N219.A, Q223.A, S224.A, L225.A, K229.A, F230.A, I35.A |
|  | PN4 | R218.A, N219.A, S222.A, R228.A, Q319.A, E316.A, I176.A, K178.A |
|  | PN5 | R307.A, Y310.A, V232.A, L314.A, E315.A, S226.A, D94.A, E233.A, L225.A |
| Phosphoprotein | PP1 | T562.B, K559.B, V556.B, N561.C, T562.C, T566.C, E568.C, I567.C |
|  | PP2 | L517.C, E514.C, V516.C, N522.C, D482.B |
| Fusion protein | PF2 | V39.B, Y30.B, H29.B, Y432.B, L433.B, N380.B, K40.B |
| Matrix protein | PM1 | E195.A, H238.A, P332.A, Q328.A, L207.A, M236.A, D304.A, M188.A |
|  | PM2 | F151.A, K143.A, W141.A, Y62.A, L181.A, Y187.A, M188.A, L274.A, D304.A |
|  | PM3 | L312.A, W314.A, L309.A, F235.A, D213.A, M236.A, F266.A |

Supporting Table 9- List of the ranks and energy values of the small drug like molecules that were predicted in the top 100 by both DOCK6.8 and Autodock4. RMSD_1 – RMSD_5 are the RMSDs of the 5 best Autodock4 poses with the best scoring Dock6.8 pose. The least RMSD is depicted in bold. Cells highlighted in yellow have RMSDs better than 0.15 nm. Pocket number indicates pockets from Autodock. Some of the Autodock pockets have been subdivided by DOCK, which indicates the subsections in each pocket.

| **Protein name** | **PDB ID** | **Pocket Number** | **Number of selected molecules** | **ZINC ID** | **Rank in DOCK** | **Rank in Autodock** | **Energy in DOCK** | **Energy in Autodock** | **RMSD_1 (nm)** | **RMSD_2(nm)** | **RMSD_3(nm)** | **RMSD_4(nm)** | **RMSD_5(nm)** |
| --- | --- | --- | --- | --- | --- | --- | --- | --- | --- | --- | --- | --- | --- |
| Glycoprotein | 3D11 | PG1 | 1 | ZINC63411510 | 10 | 60 | -58.15693 | -8.73 | 0.8 | **0.797** | 0.809 | 0.807 | 0.797 |
| Nucleoprotein | 4CO6 | PN1 | 3 | ZINC42750806 | 48 | 50 | -36.75411 | -8.41 | 0.432 | 0.397 | 0.648 | 0.263 | **0.251** |
|  |  | PN1 |  | ZINC02511792 | 62 | 62 | -36.45575 | -8.33 | 0.333 | 0.336 | 0.332 | **0.32** | 0.334 |
|  |  | PN1 |  | ZINC34083937 | 52 | 73 | -36.60975 | -8.27 | 0.64 | 0.637 | 0.64 | **0.601** | 0.623 |
|  |  | PN2 | 8 | ZINC94258465 | 24 | 96 | -36.12953 | -7.08 | 0.702 | **0.693** | 0.707 | 0.706 | 0.705 |
|  |  | PN2 |  | ZINC94258558 | 33 | 79 | -35.80085 | -7.17 | 0.081 | **0.074** | 0.074 | 0.096 | 0.076 |
|  |  | PN2 |  | ZINC86657759 | 64 | 54 | -34.88816 | -7.31 | **0.505** | 0.509 | 0.518 | 0.506 | 0.508 |
|  |  | PN2 |  | ZINC73641145 | 6 | 28 | -37.48719 | -7.6 | 0.145 | 0.148 | 0.261 | 0.149 | **0.142** |
|  |  | PN2 |  | ZINC95022396 | 98 | 42 | -34.33191 | -7.43 | 0.672 | 0.674 | 0.688 | 0.68 | **0.661** |
|  |  | PN2 |  | ZINC77262630 | 4 | 50 | -38.61404 | -7.35 | 0.275 | **0.187** | 0.232 | 0.221 | 0.255 |
|  |  | PN2 |  | ZINC72264974 | 91 | 47 | -34.37937 | -7.36 | 1.115 | 1.094 | **1.085** | 1.103 | 1.171 |
|  |  | PN2 |  | ZINC04580552 | 10 | 19 | -36.85627 | -7.79 | 0.375 | 0.376 | 0.286 | **0.272** | 0.442 |
|  |  | PN2 | 5 | ZINC73641145 | 61 | 28 | -36.67706 | -7.6 | 1.368 | 1.373 | **1.351** | 1.356 | 1.369 |
|  |  | PN2 |  | ZINC72129411 | 32 | 44 | -37.25832 | -7.41 | **1.68** | 1.693 | 1.696 | 1.686 | 1.682 |
|  |  | PN2 |  | ZINC72107957 | 87 | 8 | -36.21572 | -8.16 | **1.322** | 1.401 | 1.418 | 1.538 | 1.529 |
|  |  | PN2 |  | ZINC77262630 | 9 | 49 | -38.72784 | -7.35 | **1.378** | 1.431 | 1.442 | 1.432 | 1.418 |
|  |  | PN2 |  | ZINC94937158 | 90 | 60 | -36.05951 | -7.28 | 1.436 | 1.409 | **1.393** | 1.404 | 1.401 |
|  |  | PN4 | 10 | ZINC16932105 | 14 | 32 | -40.61285 | -9.07 | 0.917 | 0.917 | 0.92 | 0.533 | **0.524** |
|  |  | PN4 |  | ZINC12362922 | 10 | 25 | -41.36608 | -9.15 | 0.785 | 0.769 | **0.139** | 0.147 | 0.142 |
|  |  | PN4 |  | ZINC92722391 | 45 | 82 | -38.74677 | -8.61 | 0.408 | **0.394** | 0.411 | 0.497 | 0.407 |
|  |  | PN4 |  | ZINC00149964 | 57 | 29 | -38.08893 | -9.1 | 0.6 | 0.595 | 0.584 | 0.605 | **0.577** |
|  |  | PN4 |  | ZINC06361369 | 81 | 24 | -37.51473 | -9.18 | 0.837 | 0.724 | 0.781 | **0.718** | 0.842 |
|  |  | PN4 |  | ZINC02819777 | 65 | 5 | -37.90257 | -9.73 | 0.685 | 0.701 | **0.65** | 0.679 | 0.664 |
|  |  | PN4 |  | ZINC04829362 | 21 | 97 | -40.30071 | -8.53 | **0.085** | 0.121 | 0.119 | 0.119 | 0.115 |
|  |  | PN4 |  | ZINC04085190 | 39 | 33 | -38.94919 | -9.01 | **0.467** | 0.492 | 0.48 | 0.476 | 0.481 |
|  |  | PN4 |  | ZINC00814199 | 8 | 64 | -41.43409 | -8.77 | 0.553 | 0.51 | 0.514 | **0.508** | 0.526 |
|  |  | PN4 |  | ZINC92179996 | 24 | 2 | -39.96397 | -9.82 | 0.637 | 0.64 | 0.638 | 0.613 | **0.599** |
|  |  | PN5 | 4 | ZINC49587767 | 57 | 81 | -36.72681 | -6.59 | 0.655 | **0.592** | 0.631 | 0.67 | 0.62 |
|  |  | PN5 |  | ZINC04334885 | 21 | 35 | -37.89314 | -6.83 | 0.744 | 0.742 | 0.736 | 0.762 | **0.732** |
|  |  | PN5 |  | ZINC72107957 | 87 | 6 | -36.21572 | -7.65 | 0.664 | 0.66 | 0.588 | **0.506** | 0.524 |
|  |  | PN5 |  | ZINC73641145 | 61 | 52 | -36.67706 | -6.73 | 1.32 | 1.444 | 1.372 | 0.804 | **0.741** |
| Phosphoprotein | 4N5B | PP1 | 8 | ZINC85650631 | 79 | 34 | -34.26032 | -6.71 | 0.168 | **0.164** | 0.233 | 0.232 | 0.167 |
|  |  | PP1 |  | ZINC94927184 | 19 | 85 | -35.80973 | -6.55 | **0.176** | 0.19 | 0.177 | 0.395 | 0.385 |
|  |  | PP1 |  | ZINC95384460 | 83 | 49 | -34.14421 | -6.65 | 0.327 | 0.333 | 0.33 | **0.321** | 0.326 |
|  |  | PP1 |  | ZINC72462705 | 1 | 90 | -39.38969 | -6.54 | 0.195 | 0.167 | 0.13 | 0.178 | **0.121** |
|  |  | PP1 |  | ZINC86098248 | 93 | 65 | -34.07728 | -6.6 | **0.105** | 0.108 | 0.109 | 0.108 | 0.107 |
|  |  | PP1 |  | ZINC67884980 | 36 | 23 | -34.97003 | -6.79 | 0.204 | **0.195** | 0.197 | 0.228 | 0.227 |
|  |  | PP1 |  | ZINC77285117 | 38 | 41 | -34.88396 | -6.69 | **0.144** | 0.174 | 0.172 | 0.169 | 0.148 |
|  |  | PP1 |  | ZINC95022396 | 52 | 89 | -34.62392 | -6.54 | 0.261 | 0.264 | **0.256** | 0.261 | 0.266 |
|  |  | PP1 | 6 | ZINC20534353 | 37 | 75 | -34.05145 | -6.57 | 0.31 | 0.305 | 0.301 | 0.303 | **0.282** |
|  |  | PP1 |  | ZINC77379208 | 66 | 16 | -33.37782 | -6.87 | **0.191** | 0.215 | 0.214 | 0.2 | 0.197 |
|  |  | PP1 |  | ZINC94927184 | 20 | 83 | -35.13292 | -6.55 | 0.589 | 0.564 | 0.588 | 0.594 | **0.537** |
|  |  | PP1 |  | ZINC65405061 | 40 | 55 | -33.95987 | -6.62 | **0.236** | 0.239 | 0.239 | 0.24 | 0.242 |
|  |  | PP1 |  | ZINC72462705 | 9 | 89 | -36.10227 | -6.54 | **0.14** | 0.227 | 0.2 | 0.229 | 0.198 |
|  |  | PP1 |  | ZINC77285117 | 12 | 39 | -35.83965 | -6.69 | 0.177 | 0.136 | 0.152 | **0.13** | 0.18 |
|  |  | PP1 | 2 | ZINC72462705 | 10 | 89 | -62.96717 | -6.54 | 7.624 | **7.591** | 7.638 | 7.599 | 7.632 |
|  |  | PP1 |  | ZINC94927184 | 7 | 84 | -64.35318 | -6.55 | 7.708 | 7.709 | 7.716 | 7.587 | **7.584** |
|  |  | PP2 | 14 | ZINC04722076 | 5 | 30 | -68.47876 | -9.91 | **0.515** | **0.515** | **0.515** | **0.515** | **0.515** |
|  |  | PP2 |  | ZINC71260677 | 70 | 34 | -56.16101 | -9.77 | 0.218 | 0.217 | 0.213 | **0.208** | 0.209 |
|  |  | PP2 |  | ZINC94927184 | 7 | 49 | -64.29467 | -9.54 | **2.032** | 2.055 | 2.414 | 2.317 | 2.423 |
|  |  | PP2 |  | ZINC67895025 | 62 | 92 | -56.83767 | -9.25 | **0.66** | 0.672 | 0.664 | **0.66** | **0.66** |
|  |  | PP2 |  | ZINC19362297 | 74 | 67 | -55.84508 | -9.41 | 0.301 | **0.299** | **0.299** | 0.314 | 0.299 |
|  |  | PP2 |  | ZINC01584645 | 10 | 35 | -62.48652 | -9.76 | **0.943** | 1.031 | 1.027 | 1.014 | 1.073 |
|  |  | PP2 |  | ZINC86094832 | 29 | 73 | -59.68003 | -9.38 | 0.507 | **0.346** | 0.369 | 0.658 | 0.523 |
|  |  | PP2 |  | ZINC86095599 | 34 | 97 | -59.0738 | -9.21 | **0.118** | 0.165 | 0.155 | 0.133 | 0.133 |
|  |  | PP2 |  | ZINC92209154 | 35 | 29 | -59.0716 | -9.92 | 0.632 | 0.638 | **0.62** | 0.748 | 0.787 |
|  |  | PP2 |  | ZINC95221243 | 47 | 33 | -57.94121 | -9.77 | **0.949** | 0.955 | 0.961 | 0.978 | 0.957 |
|  |  | PP2 |  | ZINC91252717 | 2 | 1 | -71.46974 | -14.3 | 0.437 | 0.438 | **0.427** | 0.43 | 0.431 |
|  |  | PP2 |  | ZINC35605802 | 38 | 15 | -58.90168 | -10.29 | 0.114 | 0.115 | **0.111** | 0.116 | 0.116 |
|  |  | PP2 |  | ZINC72143751 | 91 | 27 | -55.16788 | -10.11 | **2.07** | **2.07** | 2.072 | 2.075 | 2.071 |
|  |  | PP2 |  | ZINC72462705 | 14 | 24 | -61.49333 | -10.16 | 0.955 | 0.96 | **0.866** | 0.887 | 0.89 |
| Fusion protein | 5EVM | PF2 | 5 | ZINC94725877 | 91 | 43 | -38.39184 | -8.82 | 0.397 | 0.405 | 0.4 | 0.374 | **0.311** |
|  |  | PF2 |  | ZINC72131030 | 74 | 4 | -38.72321 | -9.52 | **0.45** | 0.452 | 0.473 | 0.471 | 0.457 |
|  |  | PF2 |  | ZINC93518195 | 68 | 92 | -38.83024 | -8.5 | 0.582 | 0.582 | 0.582 | 0.585 | **0.578** |
|  |  | PF2 |  | ZINC65418720 | 7 | 94 | -42.79324 | -8.49 | **0.541** | 0.566 | 0.579 | 0.567 | 0.578 |
|  |  | PF2 |  | ZINC34083754 | 65 | 35 | -38.9075 | -8.9 | 0.362 | 0.369 | 0.385 | 0.357 | **0.343** |
| Matrix protein | Monomer of modeled dimer | PM1 | 1 | ZINC02511792 | 41 | 87 | -39.76443 | -7.45 | 0.543 | 0.503 | 0.599 | 0.504 | **0.48** |
|  |  |  |  |  |  |  |  |  | 0 | 0 | 0 | 0 | 0 |
|  |  | PM1 | 1 | ZINC02819777 | 26 | 22 | -36.58277 | -7.85 | 1.689 | 1.668 | **1.664** | 1.674 | 1.683 |
|  |  | PM2 | 12 | ZINC26481080 | 53 | 25 | -45.33724 | -8.55 | 0.415 | 0.394 | 0.414 | 0.389 | **0.388** |
|  |  | PM2 |  | ZINC12362922 | 78 | 29 | -44.42041 | -8.51 | **0.183** | 0.186 | 0.185 | **0.183** | **0.183** |
|  |  | PM2 |  | ZINC00814199 | 14 | 7 | -49.45135 | -8.97 | 0.624 | 0.621 | 0.622 | **0.619** | 0.624 |
|  |  | PM2 |  | ZINC31165406 | 34 | 20 | -46.42793 | -8.61 | 0.4 | 0.392 | **0.236** | 0.381 | 0.47 |
|  |  | PM2 |  | ZINC00149964 | 31 | 16 | -47.13465 | -8.67 | 0.624 | 0.621 | 0.622 | **0.619** | 0.624 |
|  |  | PM2 |  | ZINC01725633 | 20 | 15 | -48.50071 | -8.68 | 0.409 | 0.42 | 0.204 | 0.399 | **0.193** |
|  |  | PM2 |  | ZINC16932105 | 37 | 34 | -46.15106 | -8.48 | 0.315 | **0.314** | 0.315 | 0.315 | **0.314** |
|  |  | PM2 |  | ZINC71789643 | 73 | 30 | -44.57679 | -8.51 | 0.364 | 0.352 | 0.366 | **0.341** | 0.362 |
|  |  | PM2 |  | ZINC93518353 | 97 | 100 | -43.77466 | -8.02 | 0.41 | 0.411 | **0.397** | 0.464 | 0.492 |
|  |  | PM2 |  | ZINC91497887 | 87 | 67 | -44.09179 | -8.21 | 0.192 | 0.187 | 0.19 | **0.185** | 0.2 |
|  |  | PM2 |  | ZINC02819777 | 90 | 28 | -44.02878 | -8.51 | 0.475 | 0.469 | 0.471 | **0.414** | 0.482 |
|  |  | PM2 |  | ZINC19735365 | 86 | 93 | -44.10543 | -8.04 | **0.529** | **0.529** | 0.532 | 0.531 | 0.532 |
|  |  | PM2 | 11 | ZINC26481080 | 20 | 24 | -45.25652 | -8.55 | 0.352 | 0.355 | 0.364 | **0.293** | **0.293** |
|  |  | PM2 |  | ZINC93518353 | 36 | 98 | -43.07938 | -8.02 | 0.389 | 0.387 | 0.385 | 0.447 | **0.35** |
|  |  | PM2 |  | ZINC00814199 | 27 | 7 | -44.39831 | -8.97 | 0.655 | **0.645** | 0.653 | 0.652 | 0.682 |
|  |  | PM2 |  | ZINC00149964 | 12 | 16 | -46.60623 | -8.67 | **0.452** | 0.454 | 0.459 | 0.457 | 0.457 |
|  |  | PM2 |  | ZINC02819777 | 92 | 29 | -40.61911 | -8.51 | 0.315 | 0.316 | 0.337 | 0.384 | **0.302** |
|  |  | PM2 |  | ZINC31165406 | 22 | 20 | -45.00608 | -8.61 | 0.424 | 0.42 | **0.297** | 0.398 | 0.446 |
|  |  | PM2 |  | ZINC02511792 | 68 | 67 | -41.30507 | -8.21 | **0.165** | 0.187 | 0.185 | 0.173 | 0.166 |
|  |  | PM2 |  | ZINC91497887 | 89 | 66 | -40.66805 | -8.21 | **0.247** | **0.247** | 0.254 | 0.253 | 0.252 |
|  |  | PM2 |  | ZINC00129345 | 66 | 95 | -41.371 | -8.04 | 0.423 | 0.454 | **0.41** | 0.441 | 0.42 |
|  |  | PM2 |  | ZINC01725633 | 58 | 15 | -41.75074 | -8.68 | 0.423 | 0.418 | **0.415** | 0.418 | 0.447 |
|  |  | PM2 |  | ZINC04020772 | 23 | 46 | -44.95806 | -8.37 | 0.457 | 0.461 | 0.452 | **0.449** | 0.461 |
|  |  | PM3 | 6 | ZINC49453727 | 86 | 70 | -38.79175 | -6.56 | 1.949 | **1.94** | 1.946 | 1.969 | 1.992 |
|  |  | PM3 |  | ZINC83328368 | 25 | 33 | -40.687 | -6.82 | 1.775 | 1.781 | 1.775 | **1.74** | 1.743 |
|  |  | PM3 |  | ZINC72131030 | 72 | 19 | -38.97821 | -6.9 | 1.892 | **1.89** | 1.914 | 1.893 | 1.905 |
|  |  | PM3 |  | ZINC20390482 | 60 | 94 | -39.18076 | -6.48 | 1.821 | 1.821 | 1.821 | **1.819** | 1.82 |
|  |  | PM3 |  | ZINC63781317 | 51 | 53 | -39.36716 | -6.66 | 1.839 | 1.829 | 1.829 | **1.746** | 1.796 |
|  |  | PM3 |  | ZINC20163996 | 12 | 72 | -42.23375 | -6.56 | 1.896 | 1.896 | 1.894 | **1.893** | **1.893** |
|  |  | PM3 | 6 | ZINC20163996 | 70 | 69 | -33.02703 | -6.56 | 2.471 | 2.47 | 2.466 | 2.464 | **2.463** |
|  |  | PM3 |  | ZINC05603964 | 23 | 27 | -37.15054 | -6.85 | 3.517 | 3.53 | 3.526 | 3.524 | **3.515** |
|  |  | PM3 |  | ZINC72131030 | 48 | 19 | -34.14613 | -6.9 | 1.832 | 1.837 | **1.793** | 1.827 | 1.803 |
|  |  | PM3 |  | ZINC91497887 | 9 | 43 | -39.84722 | -6.73 | 3.406 | 3.389 | 3.426 | **3.384** | 3.425 |
|  |  | PM3 |  | ZINC73736970 | 88 | 24 | -32.46295 | -6.86 | 3.397 | **3.339** | 3.412 | 3.433 | 3.427 |
|  |  | PM3 |  | ZINC02819777 | 26 | 15 | -36.58277 | -7.1 | 3.317 | 3.338 | **3.303** | 3.342 | 3.356 |

Supporting Table 10- Number of Hydrogen bonds that are formed between the selected pose for DOCK6.8 and Autodock4 with the protein. Number of common Hydrogen bonds indicates the number of hydrogen bonds that are common between the predicted poses of the ligand from Autodock4 and DOCK6.8.

| **Sr no** | **Protein Name** | **Pocket Number** | **ZINC ID** | **Number of Hydrogen bonds for DOCK6.8** | **Number of Hydrogen bonds for Autodock4** | **Number of common Hydrogen bonds** |
| --- | --- | --- | --- | --- | --- | --- |
| 1 | Nucleoprotein | PN21 | ZINC94258558 | 4 | 3 | 3 |
| 2 | Nucleoprotein | PN21 | ZINC73641145 | 7 | 7 | 4 |
| 3 | Nucleoprotein | PN4 | ZINC12362922 | 5 | 7 | 2 |
| 4 | Phosphoprotein | PP11 | ZINC72462705 | 1 | 2 | 1 |
| 5 | Phosphoprotein | PP11 | ZINC86098248 | 1 | 1 | 0 |
| 6 | Phosphoprotein | PP11 | ZINC77285117 | 1 | 2 | 1 |
| 7 | Phosphoprotein | PP12 | ZINC72462705 | 1 | 2 | 0 |
| 8 | Phosphoprotein | PP12 | ZINC77285117 | 0 | 1 | 0 |
| 9 | Phosphoprotein | PP2 | ZINC86095599 | 1 | 1 | 0 |
| 10** | Phosphoprotein | PP2 | ZINC91252717 | 3 | 2 | 0 |
| 11 | Phosphoprotein | PP2 | ZINC35605802 | 2 | 4 | 2 |

** The RMSD between DOCK and Autodock is 0.427 nm (greater than the cutoff). This entry is included as the rank for this ligand DOCK is 2 and Autodock is 1, indicating higher confidence in the prediction

Supporting Table 11 – Same drug like molecule predicted to bind different pockets of the same or different protein. The binding pocket has been mentioned in parenthesis.

| **ZINC ID** | **Protein (Binding Pocket)** |
| --- | --- |
| ZINC00149964 | N (PN4), M (PM2) |
| ZINC00814199 | N (PN4), M (PM2) |
| ZINC02511792 | N (PN1), M (PM1, PM2) |
| ZINC02819777 | N (PN4), M (PM1, PM2, PM3) |
| ZINC12362922 | N (PN4), M (PM2) |
| ZINC16932105 | N (PN4), M (PM2) |
| ZINC72107957 | N (PN2, PN4) |
| ZINC72131030 | F (PF2), M (PM3) |
| ZINC72462705 | P (PP1, PP2) |
| ZINC73641145 | N (PN2, PN5) |
| ZINC91497887 | M (PM2, PM3) |

Supporting Table 12 – The sequence variations between the 15 NiV strains. The mutations are mentioned by the residue number followed by the amino acids present in different strains.

| **Protein** | **Conservation (%)** | **Mutations** |
| --- | --- | --- |
| C | 94.0 | 25 A/V, 33 I/T, 34 K/E, 37 H/P, 39 K/R, 40 I/T, 62 M/V, 67 A/T, 83 K/R, 98 Y/H |
| F | 98.0 | 2 A/V, 6 D/N, 9 Y/C, 11 S/C, 15 I/L, 19 I/M, 42 I/V, 207 S/L, 250 I/T, 252 D/G, 273 S/G |
| G | 94.5 | 3 A/T, 5 S/N, 14 A/T, 20 I/N, 82 M/V, 89 S/G, 172 K/R, 236 K/R, 272 A/T, 274 P/S, 288 S/N, 299 T/V, 304 I/V, 325 S/N, 328 E/G, 329 S/G, 335 L/F, 339 S/N, 344 K/R/M, 384 I/V, 385 A/T, 386 K/E, 421 E/G, 424 P/S, 426 I/V, 427 I/V, 470 Q/L, 481 D/N, 498 K/T, 502 I/V, 545 I/V |
| L | 98.0 | 36 K/R, 71 D/N, 77 I/V, 94 I/T, 112 K/R, 223 T/N, 252 I/V, 533 E/D, 621 K/R, 625 Y/C, 632 S/N, 639 D/N, 640 P/S, 642 Y/N, 658 Y/H, 665 I/T, 703 K/R, 783 K/E, 890 I/V, 1154 I/L, 1157 K/R, 1181 K/R, 1262 K/R, 1494 A/V, 1551 A/S, 1577 I/V, 1645 Y/S/F, 1658 S/N, 1707 M/V, 1748 I/V, 1753 M/V, 1791 A/S, 1801 K/R, 1896 A/T, 2001 M/V, 2027 I/V, 2031 K/R, 2037 D/N, 2039 H/N, 2064 E/D, 2071 Q/H, 2159 C/R, 2216 S/N |
| M | 98.6 | 13 I/M, 26 H/N, 127 I/V, 147 S/G, 331 I/V |
| N | 97.4 | 139 S/R, 188 E/D, 211 Q/R, 345 I/M, 387 D/N, 429 I/V, 432 E/G, 457 D/N, 503 S/N, 505 K/R, 506 D/T, 508 R/G, 520 P/S, 521 A/T |
| P | 89.3 | 41 Q/R, 64 P/S, 69 D/G, 74 S/N, 105 I/T, 139 Y/H, 140 S/T, 147 D/N, 156 M/V, 179 D/N, 183 A/T, 191 I/V, 195 P/L, 196 K/R, 200 D/V, 218 K/R, 219 E/G, 223 D/G, 225 Q/E, 227 S/N, 228 K/R, 269 E/D, 274 S/R, 275 A/V, 276 S/G, 277 R/G, 280 I/N, 283 A/I/V, 285 H/R, 286 I/T, 287 I/L, 292 I/T, 295 S/N, 297 Q/K, 298 I/A, 300 D/G, 303 P/S, 304 A/T, 306 A/V, 310 R/G, 311 P/L, 319 K/E, 320 P/S, 343 Q/R, 351 L/F, 354 S/C, 363 P/L, 365 Y/H, 366 R/W, 367 S/G, 370 R/G, 372 I/R, 377 A/T, 378 K/E, 380 T/V, 381 S/N, 382 D/G, 386 T/N, 388 D/N, 389 K/R, 410 A/E, 421 P/L, 425 S/N, 449 Q/R, 452 A/V, 453 P/S, 455 A/V, 464 A/V, 467 A/V, 590 S/N, 602 I/V, 629 A/T, 635 E/G, 664 I/V, 683 D/G, 687 K/R |
| V | 78.9 | 41 Q/R, 64 P/S, 69 D/G, 74 S/N, 105 I/T, 139 Y/H, 140 S/T, 147 D/N, 156 M/V, 179 D/N, 183 A/T, 191 I/V, 195 P/L, 196 K/R, 200 D/V, 218 K/R, 219 E/G, 223 D/G, 225 Q/E, 227 S/N, 228 K/R, 269 E/D, 275 A/V, 277 R/G, 280 I/N, 283 A/I/V, 285 H/R, 286 I/T, 287 I/L, 292 I/T, 295 S/N, 297 Q/K, 298 I/A, 300 D/G, 303 P/S, 304 A/T, 306 A/V, 311 P/L, 319 K/E, 320 P/S, 343 Q/R, 351 L/F, 354 S/C, 363 P/L, 365 Y/H, 366 R/W, 367 S/G, 370 R/G, 372 I/R, 377 A/T, 380 T/V, 381 S/N, 382 D/G, 386 T/N, 388 D/N, 389 K/R, 408 H/T, 409 R/D, 410 R/E, 411 K/E, 412 Y/I, 413 P/S, 414 I/S, 415 A/C, 416 W/G, 417 T/D, 418 E/G, 419 K/N, 424 P/E/L, 425 E/G, 426 S/W, 427 K/C, 428 S/N, 429 P/G, 431 C/T, 432 S/R, 433 H/R, 434 I/V, 435 R/T, 436 P/G, 437 S/L, 439 P/R, 440 Y/R, 443 C/G, 444 Q/K, 445 S/C, 446 G/V, 447 E/N, 448 A/C, 452 Q/R/, 458 E/V, 459 C/V, 460 K/F, 461 H/E, 462 C/T, 463 D/G |
| W | 79.3 | 41 Q/R, 64 P/S, 69 D/G, 74 S/N, 105 I/T, 139 Y/H, 140 S/T, 147 D/N, 156 M/V, 179 D/N, 183 A/T, 191 I/V, 195 P/L, 196 K/R, 200 D/V, 218 K/R, 219 E/G, 223 D/G, 225 Q/E, 227 S/N, 228 K/R, 269 E/D, 275 A/V, 277 R/G, 280 I/N, 283 A/I/V, 285 H/R, 286 I/T, 287 I/L, 292 I/T, 295 S/N, 297 Q/K, 298 I/A, 300 D/G, 303 P/S, 304 A/T, 306 A/V, 311 P/L, 319 K/E, 320 P/S, 343 Q/R, 351 L/F, 354 S/C, 363 P/L, 365 Y/H, 366 R/W, 367 S/G, 370 R/G, 372 I/R, 377 A/T, 380 T/V, 381 S/N, 382 D/G, 386 T/N, 388 D/N, 389 K/R, 408 A/T, 409 Q/D, 410 E/T, 411 K/R, 412 Y/N, 413 I/P, 414 H/S, 416 A/L, 418 R/T, 419 K/E, 420 T/N, 421 C/V, 422 P/L, 424 S/R, 425 K/R, 426 S/V/N, 427 G/V, 428 A/Q, 429 P/T, 430 R/G, 431 H/M, 432 F/V, 433 R/E, 434 D/G, 435 H/S, 438 Y/T, 439 Q/K, 440 K/E, 441 A/G, 442 K/R 445 S/N, 446 A/M, 447 R/E, 448 R/N, 449 M/V, 450 Q/S/R, 451 L/N |

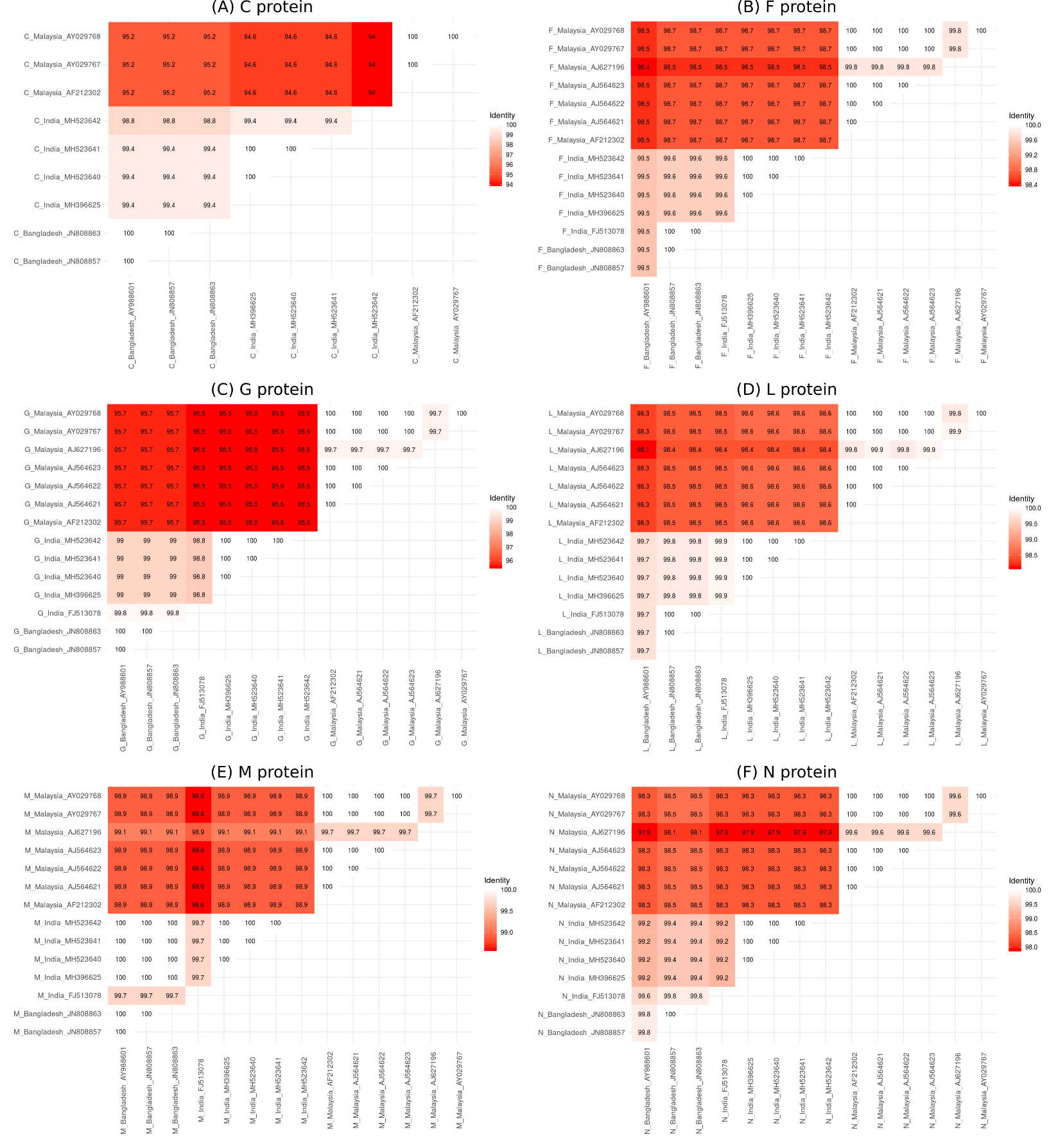

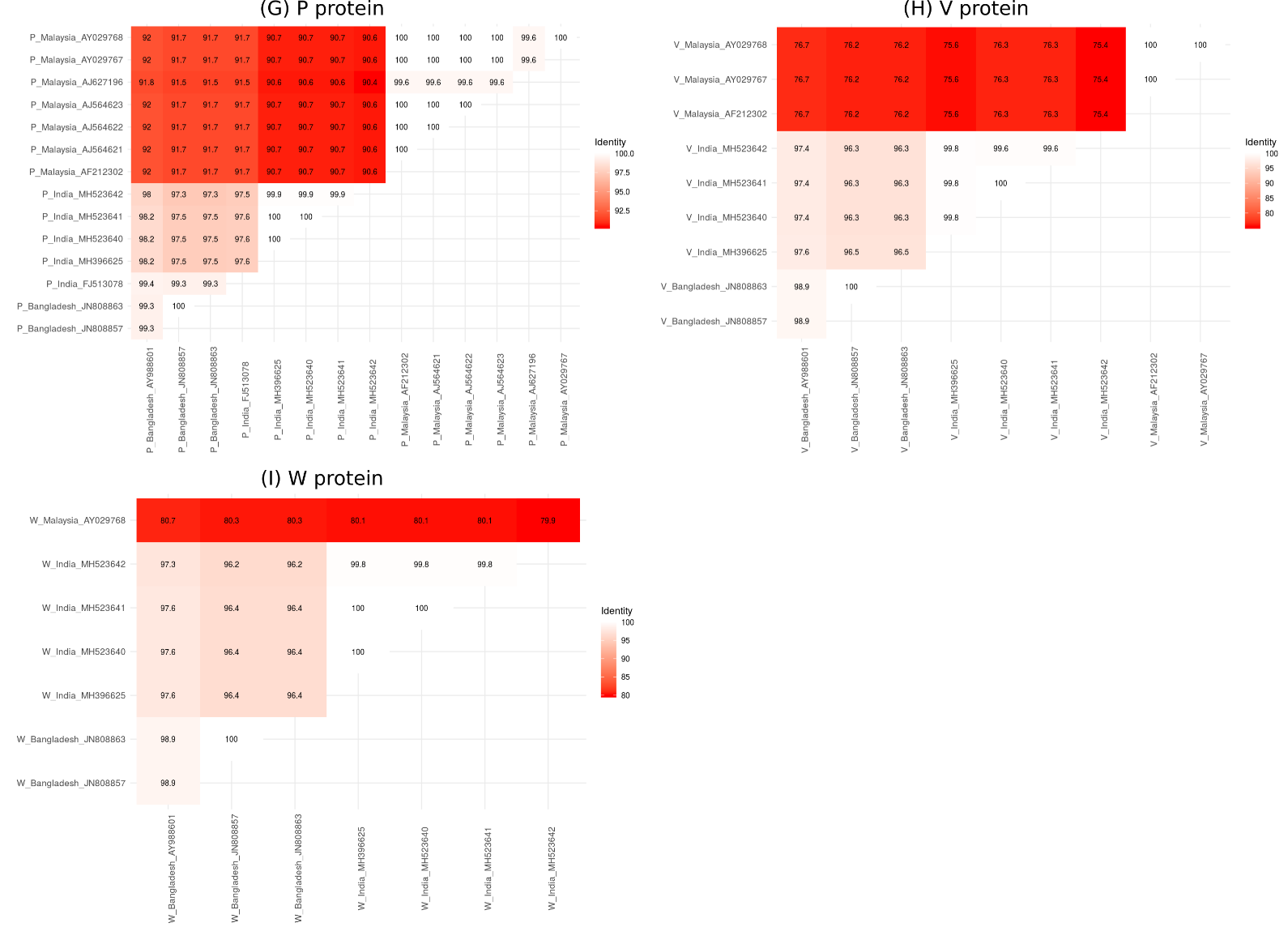

Supporting Figure 6 – Heatmap showing the sequence conservation between the different strains of NiV for (A) C protein (B) F protein (C) G protein (D) L protein (E) M protein (F) N protein (G) P protein (H) V protein (I) W protein. The color gradient represents sequence conservation where white indicates 100% conservation and redder shades indicate lesser sequence conservation. The labelling convention is Protein_Country_Genome-accession code.
